# A fungal member of the *Arabidopsis thaliana* phyllosphere antagonizes *Albugo laibachii* via a secreted lysozyme

**DOI:** 10.1101/2020.04.20.051367

**Authors:** Katharina Eitzen, Priyamedha Sengupta, Samuel Kroll, Eric Kemen, Gunther Doehlemann

## Abstract

Plants are not only challenged by pathogenic organisms, but also colonized by commensal microbes. The network of interactions these microbes establish with their host and amongst each other is suggested to contribute to the immune responses of plants against pathogens. In wild *Arabidopsis thaliana* populations, the oomycete pathogen *Albugo laibachii* has been shown to play an influential role in structuring the leaf phyllosphere. We show that the epiphytic yeast *Moesziomyces bullatus* ex *Albugo* on *Arabidopsis,* a close relative of pathogenic smut fungi, is an antagonistic member of the *A. thaliana* phyllosphere, which reduces infection of *A. thaliana* by *A. laibachii*. Combination of transcriptome analysis, reverse genetics and protein characterization identified a GH25 hydrolase with lysozyme activity as the major effector of this microbial antagonism. Our findings broaden the understanding of microbial interactions within the phyllosphere, provide insights into the evolution of epiphytic basidiomycete yeasts and pave the way for the development of novel biocontrol strategies.

## Introduction

Plants are colonized by a wide range of microorganisms. While some microbes enter the plant and establish endophytic interactions with a broad range of outcomes from beneficial to pathogenic, plant surfaces harbor a large variety of microbial organisms. Recent research has focused largely on the importance of the rhizosphere microbiota in nutrient acquisition, protection from pathogens, and boosting overall plant growth and development (1–3). However, the above ground parts of the plant including the phyllosphere are colonized by diverse groups of microbes that also assist in plant protection and immunity (4,5). The environment has a major impact on the microbial communities of the leaf surface, ultimately influencing their interactions with the host (6).

Scale-free network analysis was performed with the leaf microbial population of *Arabidopsis thaliana* (7). The majority of the interactions between kingdoms, e.g. fungi and bacteria, were found to be negative, consistent with the fact that rather the antagonistic interactions stabilize a microbial community (8). Phyllosphere network analysis of *A. thaliana* identified a small number of microbes as “hub” organisms, i.e. influential microbes which have severe effects on the community structure. The major hub microbe in the *A. thaliana* phyllosphere is the oomycete *Albugo laibachii*, which is a pathogenic symbiont biotrophic of *Arabidopsis* (7). This pathogen has been shown to significantly reduce the bacterial diversity of epiphytic and endophytic leaf habitats. Since bacteria generally comprise a large proportion of the phyllosphere microbiome (9), phylogenetic profiling of *A. thaliana* was also directed towards identifying a small group of bacteria that frequently colonize *A. thaliana* leaves. The analysis helped to develop a synthetic community of bacteria for experiments in gnotobiotic plants.

Besides bacteria and oomycetes, the microbiota of the *A. thaliana* leaf also comprises a broad range of fungi. Among those fungi, basidiomycete yeasts are frequently found and the most frequent ones are the epiphytic basidiomycete genus *Dioszegia* (7), as well as an anamorphic yeast associated with *A. laibachii* infection, classified as *Pseudozyma. sp.* and belonging to Ustilaginales. This order includes many pathogens of important crop plants, for example corn smut and loose smut of oats, barley and wheat are caused by *Ustilago maydis*, *U. avenae*, *U. nuda* and *U. tritici*, respectively. Generally, pathogenic development of smut fungi is linked with sexual recombination and plant infection is only initiated upon mating when two haploid sporidia form a dikaryotic filament (10). Ustilaginales *Pseudozyma* sp. yeasts, however, are not known to be pathogenic. While they are found in anamorphic stage, they epiphytically colonize a wide range of habitats via where an infrequent sexual recombination might occur (11). Phylogenetic reconstruction (12) showed that the smut pathogen of millet, *Moesziomyces bullatus* and four species of *Pseudozyma*, namely *P. antarctica*, *P. aphidis*, *P. parantarctica* and *P. rugulosa* form a monophyletic group. The latter does represent anamorphic and culturable stages of *M. bullatus* and, hence, can be grouped to this genus. *Moesziomyces* strains have been reported in a number of cases to act as microbial antagonists. A strain formerly classified as *Pseudozyma aphidis* (now *Moesziomyces bullatus*) inhibited *Xanthomonas campestris* pv. *vesicatoria*, *X. campestris* pv. *campestris*, *Pseudomonas syringae* pv. *tomato*, *Clavibacter michiganensis*, *Erwinia amylovora*, and *Agrobacterium tumefaciens in-vitro* and also led to the activation of induced defense responses in tomato against the pathogen (13). It was reported that *P. aphidis* can parasitize the hyphae and spores of *Podosphaera xanthii* (14). *Pseudozyma churashimaensis* was reported to induce systemic defense in pepper plants against *X. axonopodis*, Cucumber mosaic virus, Pepper mottle virus, Pepper mild mottle virus, and broad bean wilt virus (15).

In the present study, we explored the antagonistic potential of an anamorphic Ustilaginales yeast within the leaf microbial community of *A. thaliana*. We show that *Moesziomyces bullatus* ex *Albugo* on *Arabidopsis* (which will be referred to as MbA, from further on in this paper) prevents infection by the oomycete pathogen *A. laibachii* and identified fungal candidate genes that were upregulated in the presence of *A. laibachii*, when both the microbes were co-inoculated in the host plant. A knockout mutant of one of the candidates, which belongs to the glycoside hydrolase – family 25 (GH25), was found to lose its antagonistic abilities towards *A. laibachii,* providing mechanistic insights into fungal-oomycete antagonism within the phyllosphere microbiota. Functional characterization of GH25 will be an important step towards establishing *MbA* as a suitable biocontrol agent.

## Results

In a previous study we isolated a basidiomycetous yeast from *Arabidopsis thaliana* leaves infected with the causal agent of white rust, *Albugo laibachii* (7). This yeast was tightly associated with *A. laibachii* spore propagation. Even after years of subculturing in the lab and re-inoculation of plants with frozen stocks of *A. laibachii* isolate Nc14, this yeast remained highly abundant in spore isolates. Phylogenetic analyses based on fungal ITS-sequencing identified the yeast as *Pseudozyma sp*. Those yeasts can be found across the family of Ustilaginaceae, being closely related to pathogens of monocots like maize, barley, sugarcane or sorghum (Figure 1A and (16)). Microscopic analyses verified the morphological similarity between the putative *Pseudozyma sp*. and the Ustilaginaceous pathogen *Ustilago maydis*, the causal agent of corn smut (Figure 1B; (16)). Based on phylogenetic similarity to the pathogenic smut *Moesziomyces bullatus* which infects millet, several anamorphic *Pseudozyma* isolates were suggested to be renamed and grouped to *M. bullatus* (12). Since the *Pseudozyma sp*. that was isolated from *A. laibachii* spores groups into the same cluster, we classified this newly identified species as *MbA (Moesziomyces bullatus* ex *Albugo* on *Arabidopsis*).

**Figure 1:**
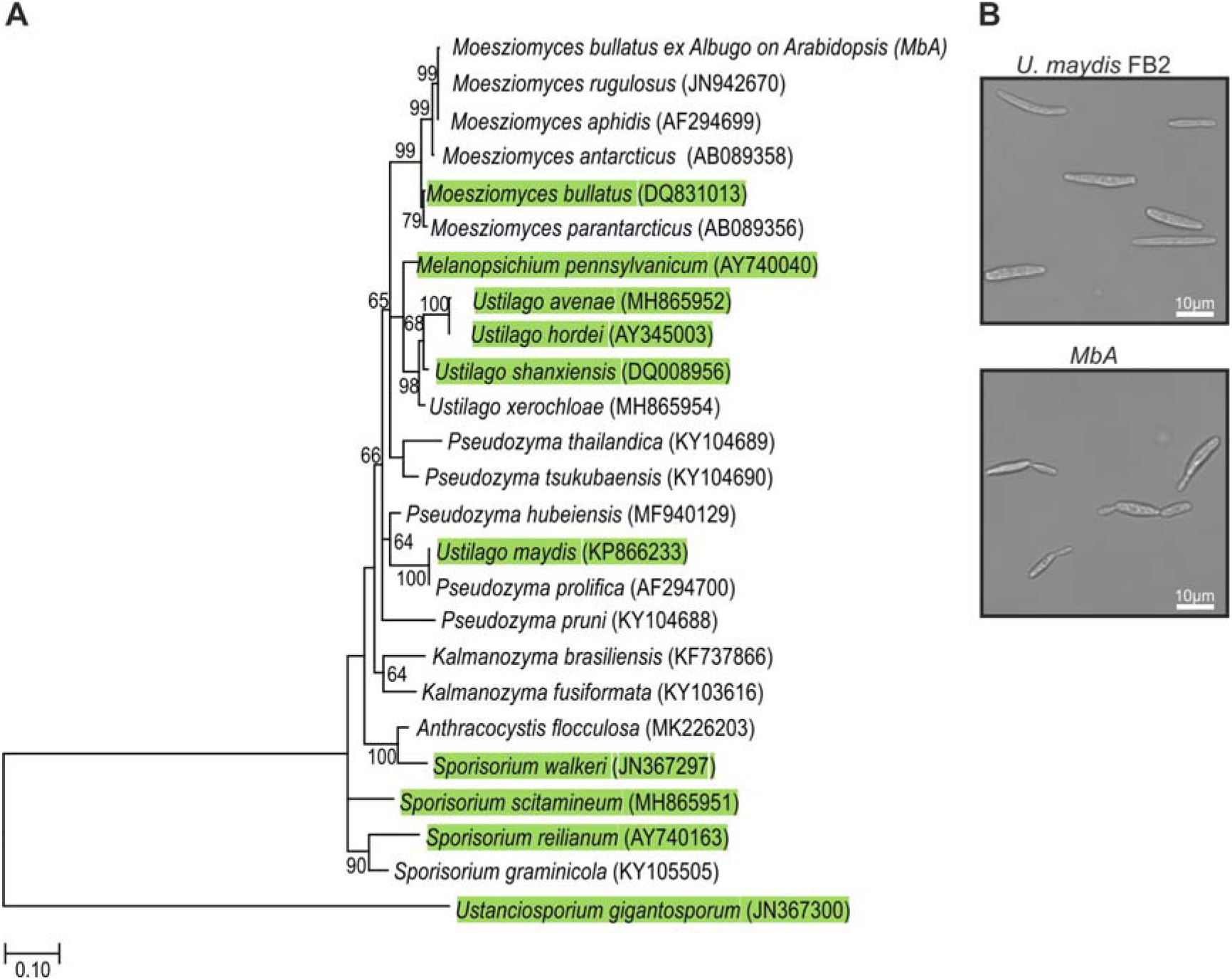
Position of *MbA* in the family of Ustilaginaceae. (A) molecular phylogenetic analysis by maximum likelihood method based on fungal ITS sequences and showing grouping of *MbA* with the millet pathogen *M. bullatus*. Pathogenic filamentous smuts (green) and anamorphic smut yeasts can be found across the phylogenetic tree. (B) Comparison of the morphology of *MbA* (bottom) and *U. maydis* (top) haploid cells grown in YEPSl_ight_ medium to OD_600_0.6.

Based on the identification of *MbA* as heaving a significant effect on bacterial diversity in the *Arabidopsis* phyllosphere, we tested its interaction with 30 bacterial strains from 17 different species of a synthetic bacterial community (SynCom, Table S1) of *Arabidopsis* leaves in one-to-one plate assays. This experiment identified seven strains being inhibited by *Moesziomyces,* as indicated by halo formation after 7 days of co-cultivation (Supplementary Figure S1). Interestingly, this inhibition was not seen when the pathogenic smut fungus *U. maydis* was co-cultivated with the bacteria, indicating a specific inhibition of the bacteria by *MbA* (Supplementary Figure S1).

The primary hub microbe in the *Arabidopsis* phyllosphere was found to be the pathogenic oomycete *A. laibachii,* which was isolated in direct association with *Moesziomyces* (7). To test if both species interfere with each other, we deployed a gnotobiotic plate system and quantified *A. laibachii* infection symptoms on *Arabidopsis*. In control experiments, spray inoculation of only *A. laibachii* spores on *Arabidopsis* leaves led to about 33% infected leaves at 14 dpi (Figure 2). When the bacterial SynCom was pre-inoculated on leaves two days before *A. laibachii* spores a significant reduction of *A. laibachii* infection by about 50% was observed (Figure 2). However, if *Moesziomyces* was pre-inoculated with the bacterial SynCom, *A. laibachii* spore production was almost completely abolished. Similarly, the pre-inoculation of only *MbA* resulted in an almost complete loss of *A. laibachii* infection, independently of the presence of a bacterial community (Figure 2). The antagonistic effect of *MbA* towards *A. laibachii* was further confirmed using Trypan blue staining of *A. laibachii* infected *A. thaliana* leaves. *A. laibachii* forms long, branching filaments on *Arabidopsis* leaves at 15 dpi. Contrary, in presence of *MbA*, we observed mostly zoospores forming either no or very short hyphae, while further colonization of the leaf with long, branching was not observed (Supplementary figure S2B). Together, of findings demonstrates that *MbA* holds a strong antagonistic activity towards *A. laibachii*, resulting in efficient biocontrol of pathogen infection. Thus, *MbA* is an important member of the *A. thaliana* phyllosphere microbial community. However, despite, several reports of the basidiomycete yeasts acting as antagonists, genomic analysis of the said group is rather limited. We therefore sequenced the genome of *MbA* and established molecular tools allowing functional genetic approaches.

**Figure 2:**
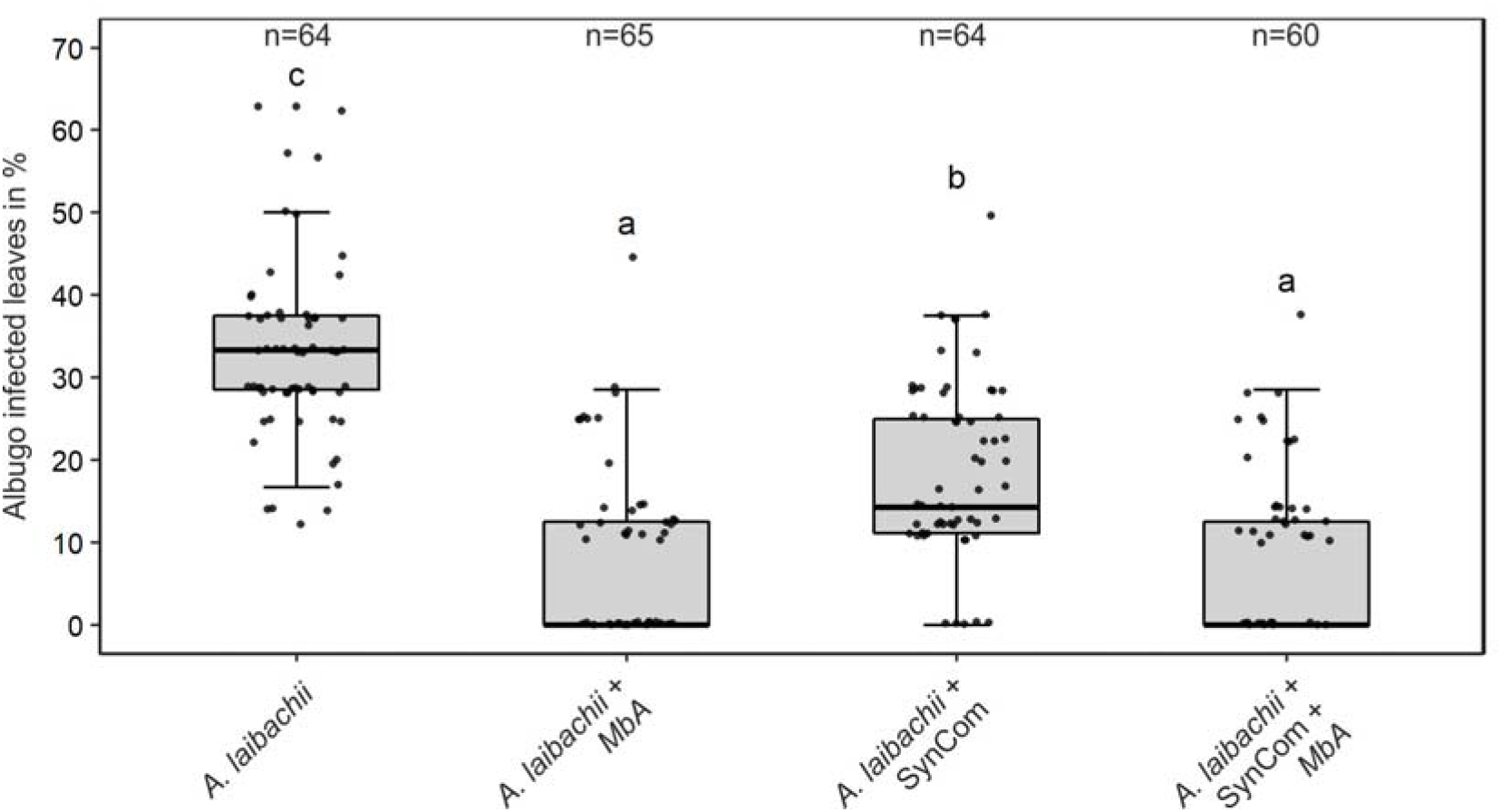
Infection assay of *A. laibachii* on *A. thaliana*. Addition of a bacterial SynCom reduces the infection symptoms of *A. laibachii* at 14dpi. Those symptoms can be almost abolished by spraying *MbA* to the plant, independently of the presence of the bacterial community. Infections were performed in six individual replicates with 12 technical replicates. N indicates the number of infected plants that were scored for symptoms. An analysis of variance (ANOVA) model was used for pairwise comparison of the conditions, with Tukey’s HSD test to determine differences among them. Different letters indicate significant differences (P values < 0.05).

### The genome of *MbA*

Genome sequence of *MbA* was analyzed by Single Molecule Real-Time sequencing (Pacific Biosciences, Menlo Park, CA), which lead to 69674 mapped reads with an accuracy of 87.3% and 8596bp sub-read length. Sequence assembly using the HGAP-pipeline (Pacific Biosciences) resulted in 31 Contigs with a N_50_Contig Length of 705kb. The total length of all contigs results in a predicted genome size of 18.3Mb (Table 1). Gene prediction for the *MbA* genome with Augustus (17) identified 6653 protein coding genes, of which 559 carry a secretion signal. Out of these 559, 380 are predicted to be secreted extracellularly (i.e. they do not carry membrane domains or cell-wall anchors) (Table 1). The small genome size and high number of coding genes results in a highly compact genome structure with only small intergenic regions. These are features similarly found in several pathogenic smut fungi such as *U. maydis* and *S. reilianum* (Table 1). Remarkably, both *MbA* and *Anthracocystis flocculosa,* which is another anamorphic and apathogenic yeast, show a similarly high rate of introns, while the pathogenic smut fungi have a significantly lower intron frequency (Table 1).

**Table 1:**
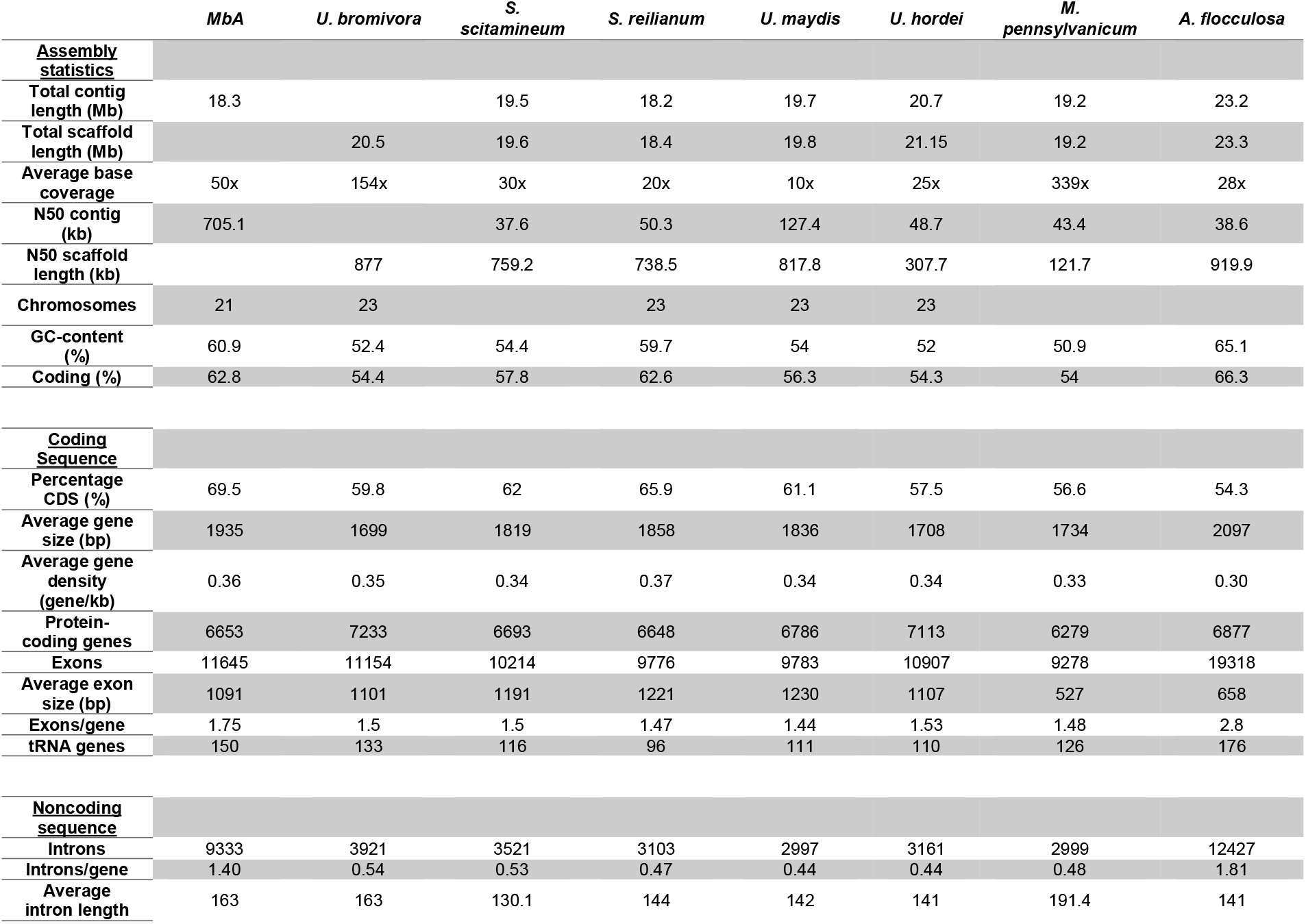
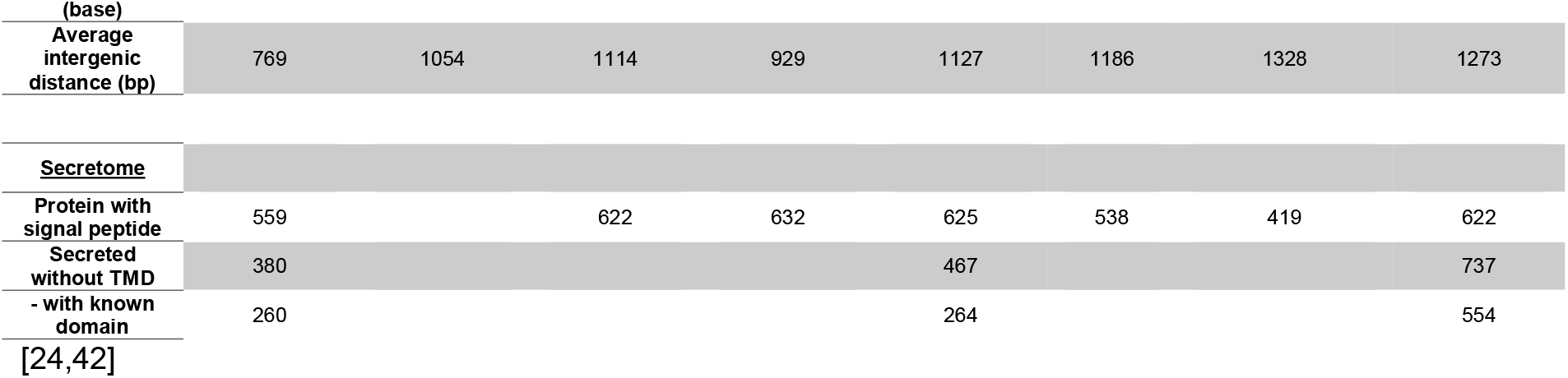
Comparison of Genomes and genomic features of known pathogenic and anamorphic Ustilaginomycetes.

To gain better insight in the genome organization of *MbA,* we compared its structure with the *U. maydis* genome, which serves as a manually annotated high-quality reference genome for smut fungi (18). Out of the 31 *MbA* contigs, 21 show telomeric structures and a high synteny to chromosomes of *U. maydis,* with three of them displaying major events of chromosomal recombination (Figure 3A). Interestingly, the *Moesziomyces* contig 2, on which also homologs to pathogenic loci like the *U. maydis* virulence cluster 2A (18) can be found, contains parts of three different *U. maydis* chromosomes (Chr. 2, 5, 20) (Supplementary figure S3). The second recombination event on contig 6 affects the *U. maydis* leaf-specific virulence factor *see1*, which is required for tumor formation (19). This recombination event is also found in the genome of the maize head smut *S. reilianum*, wherein the *U. maydis* chromosomes 5 and 20 recombined in the promoter region of the *see1* gene (Figure 3B). In this respect it should be noted that *S. reilianum*, although infecting the same host, does not produce leaf tumors as *U. maydis* does (20).

**Figure 3:**
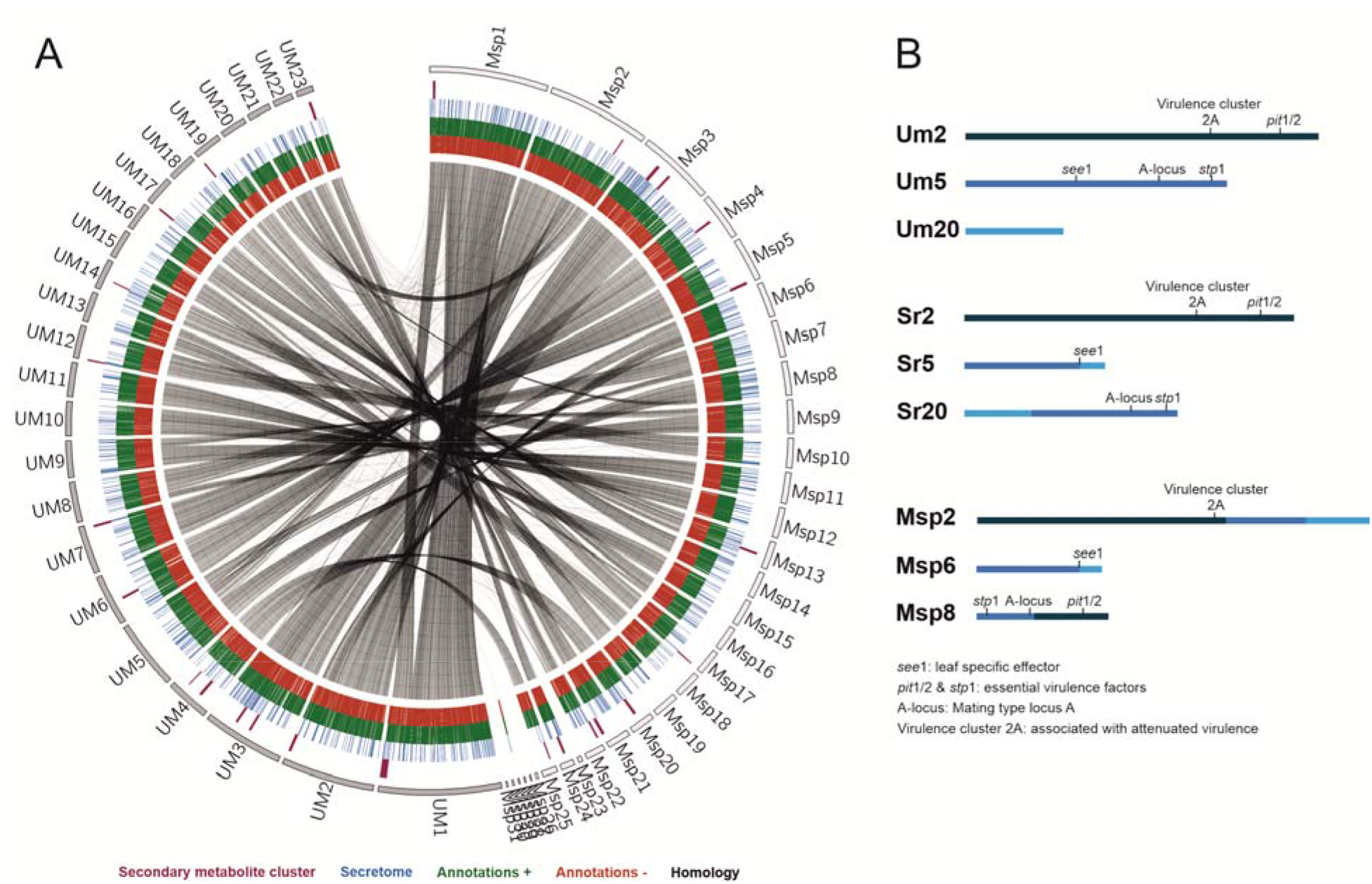
Circos comparison of *MbA* and *U. maydis* chromosome structure (A). We highlighted potential secondary metabolite clusters, secreted proteins and gene predictions on both strands (+/−). (B) Homology based comparisons identified three chromosomal recombination events, which affects the *MbA* contigs 2, 6 & 8.

Also the third major recombination event, affecting *MbA* contig 8, changes the genomic context genes encoding essential virulence factors in *U. maydis* (*stp1* & *pit1/2*), as well as the A mating type locus, which is important for pheromone perception and recognition of mating partners (21). Based on the strong antibiotic activities of *MbA,* we mined the genome of *MbA* for the presence of secondary metabolite gene clusters. Using AntiSMASH, we were able to predict 13 of such clusters, of which three can be assigned to terpene synthesis, three contain non-ribosomal peptide synthases and one cluster has a polyketide synthetase as backbone genes (Supplementary Figure S4A). Interestingly, the secondary metabolite cluster that is involved in the production of the antimicrobial metabolite ustilagic acid in other Ustilaginomycetes, is absent in *MbA* (Supplementary Figure S4B). On the contrary, we could identify three *MbA* specific metabolite clusters which could potentially be involved in the antibacterial activity of *MbA* (Supplementary Figure S4C).

A previous genome comparison of the related Ustilaginales yeast *A. flocculosa* with *U. maydis* concluded that this anamorphic strain had lost most of its effector genes, reflecting the absence of a pathogenic stage in this organism (22). In contrast, *MbA* contains 1:1 homologs of several known effectors with a known virulence function in *U. maydis* (Table 2). We previously found that *Moesziomyces* sp. possess functional homologues of the *pep1* gene, a core virulence effector of *U. maydis* (23), suggesting that such anamorphic yeasts have the potential to form infectious filamentous structures by means of sexual reproduction (11). To assess the potential virulence activity of *MbA* effector homologs, we expressed the homolog of the *U. maydis* core effector Pep1 in an *U. maydis pep1* deletion strain (SG200Δ01987). This resulted in complete restoration of *U. maydis* virulence, demonstrating that, when ectopically expressed, *MbApep1* encodes a functional effector (Supplementary Figure S5).

**Table 2:** *MbA* proteins homologous to *U. maydis* effector genes with known virulence function.

| Name | Homologue | Query cover | E-value | Identity (%) | *U. maydis* knockout phenotype | Reference |
| --- | --- | --- | --- | --- | --- | --- |
| **g1653** | UMAG_01987 (Pep1) | 82% | 3-e56 | 60.96 | complete loss of tumor formation - blocked in early stages of infection | (53) |
| **g1828** | UMAG_01829 (Afu1) | 99% | 0.0 | 71.57 | organ specific effector - reduced virulence in seedling leaves | (54) |
| **g2626** | UMAG_12197 (Cce1) | 98% | 1e-48 | 60.16 | complete loss of tumor formation - blocked in early stages of infection | (55) |
| **g2765** | UMAG_11938 (Scp2) | 100% | 1e-73 | 93.44 | Reduced in virulence | (56) |
| **g2910** | UMAG_02475 (Stp1) | 32% | 3e-42 | 60.71 | complete loss of tumor formation - blocked in early stages of infection | (57) |
| **g3652** | UMAG_02239 (See1) | 43% | 9e-11 | 54.90 | organ specific effector - reduced virulence in seedling leaves | (19) |
| **g3113** | UMAG_01375 (Pit2) | * | * | * | complete loss of tumor formation - blocked in early stages of infection | (58) |
| **g3279** | UMAG_03274 (Rsp3) | 10% | 5e-20 | 70.11 | strong attenuation of virulence – reduced tumor size and number | (59) |
| **g5296** | UMAG_05731 (Cmu1) | 98% | 3e-70 | 43.84 | Reduced virulence | (60) |
| **g6183** | UMAG_06098 (Fly1) | 100% | 0.0 | 81.85 | Reduced virulence | (61) |
| **g5835** | UMAG_05302 (Tin2) | 87% | 8e-24 | 37.81 | Minor impact on tumor formation – reduced anthocyanin biosynthesis | (24) |

A hallmark of the *U. maydis* genome structure is the presence of large clusters with effector genes, the expression of which is only induced during plant infection (18). To assess the presence of potential virulence clusters in *MbA*, we compared all *U. maydis* effector gene clusters to the *MbA* genome, based on homology. This revealed that the twelve major effector clusters of *U. maydis* are present in *MbA*. However, while many of the clustered effector genes are duplicated in pathogenic smut fungi, *MbA* carries only a single copy of each effector gene. This results in the presence of “short” versions of the *U. maydis* gene effector clusters (Supplementary Figure S6). This gets particularly obvious for the biggest and most intensively studied virulence cluster of smut fungi, the effector cluster 19A (20,24,25). In *MbA* only three out of the 24 effector genes present in *U. maydis* are conserved in this cluster (Figure 4). Interestingly, some anamorphic yeasts like *Kalmanozyma brasiliensis* and *A. flocculosa* completely lost virulence clusters, while another non-pathogenic member of the Ustilaginales, *Pseudozyma hubeiensis,* shows an almost complete set of effectors when compared to *U. maydis* (Figure 4).

**Figure 4:**
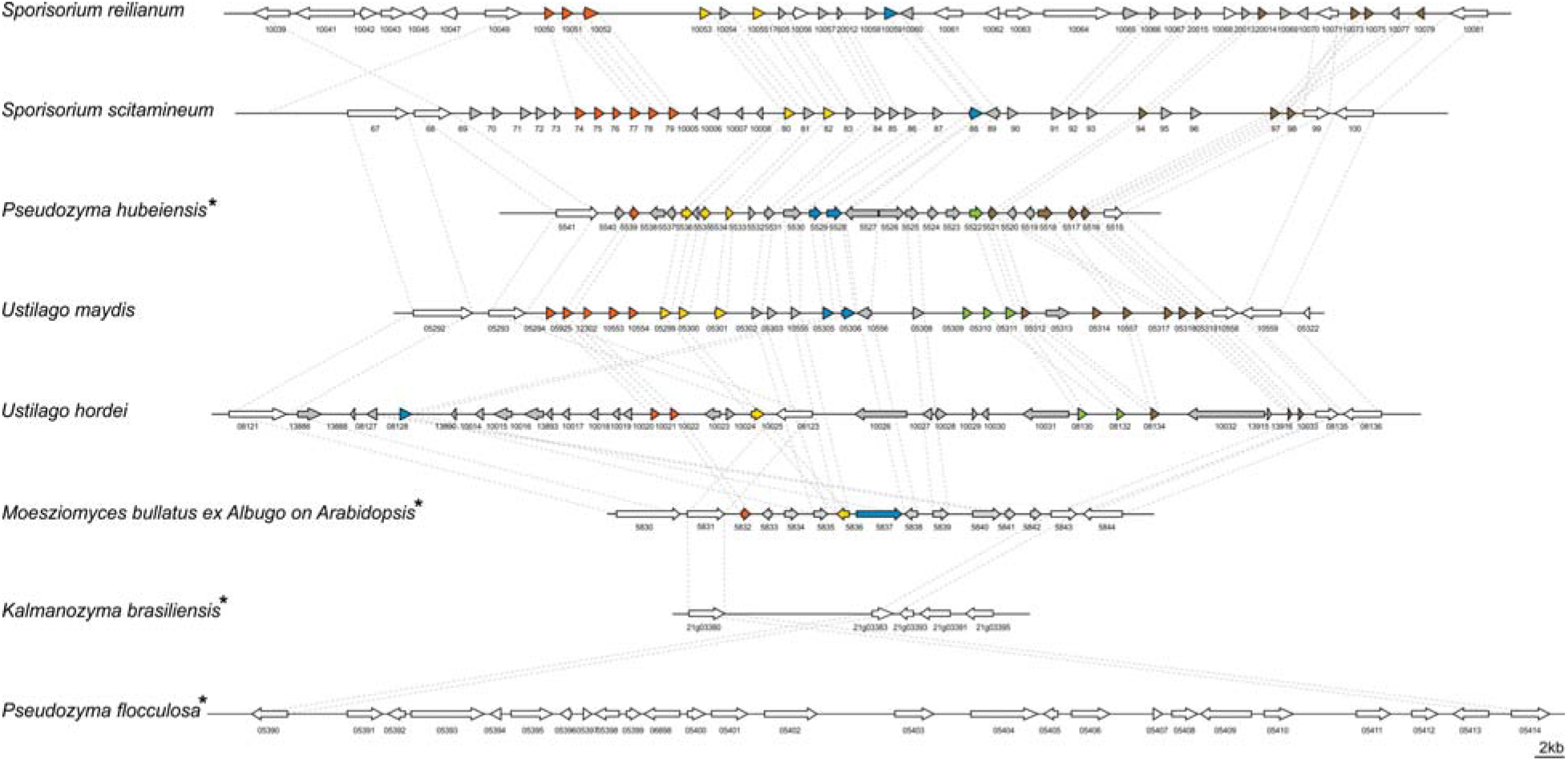
Structure of the largest virulence cluster (Cluster 19A) in pathogenic smut fungi and anamorphic smut yeasts (marked with*). Colors indicate genes with homology to each other: Related gene families are indicated in orange, yellow, blue, green and brown, whereas unique effector genes are shown in grey. Genes encoding proteins without a secretion signal are shown in white (24).

### Genetic characterization of *MbA*

To perform reverse genetics in *MbA,* we established a genetic transformation system based on protoplast preparation and PEG-mediated DNA transfer. In preliminary transformation assays, we expressed a cytosolic GFP reporter-gene under control of the constitutive *o2tef*-Promoter (Figure 5A). For the generation of knockout strains, a split marker approach was used to avoid ectopic integrations (Figure 5B). To allow generation of multiple knockouts, we used a selection marker-recycling system (pFLPexpC) which allows selection marker excision at each transformation round (26).

**Figure 5:**
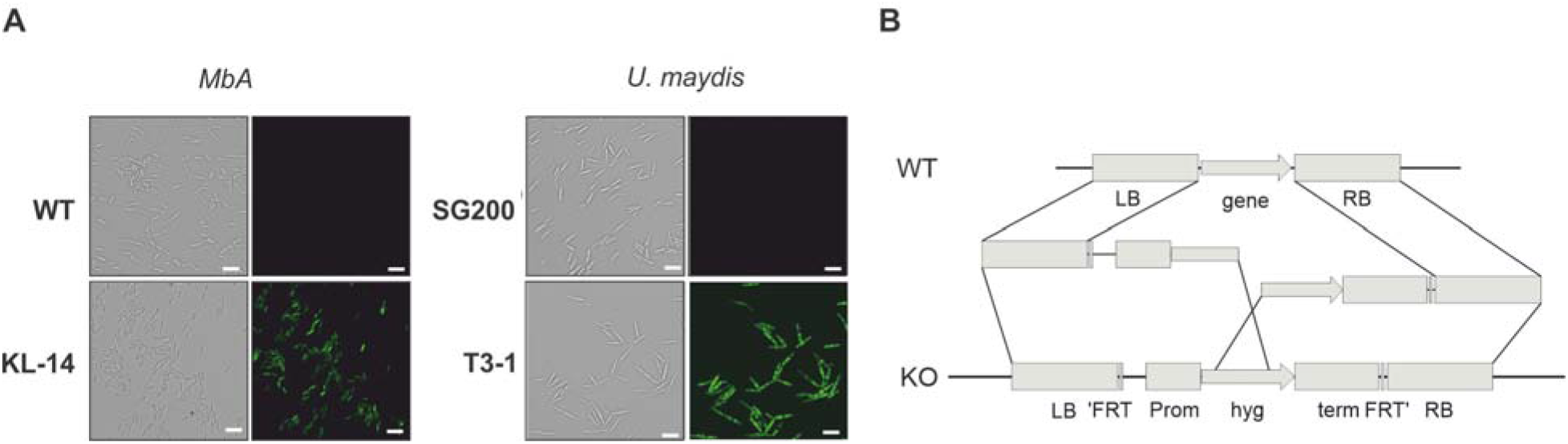
Genetic transformation of *MbA*. (A) Stable transformants that express cytosolic GFP could be obtained by generating protoplasts with Glucanex and ectopically integrating linear DNA-fragments into the genome via PEG-mediated transformation. (B) Overview of the split-marker approach that was used to generate deletion mutants via homologous recombination.

We decided to apply the transformation system to study the *MbA* mating type loci in more detail. Although phylogenetically closely related to *U. hordei,* which has a bi-polar mating system, *MbA* owns a tetrapolar mating system whereby both mating type loci are physically not linked. This situation is similar to the mating type structure in the pathogenic smut *U. maydis* (Figure 5A). The *a*-locus, which encodes a pheromone – receptor system that is required for sensing and fusion of compatible cells, is located on contig 6. The *b*-locus can be found on contig 1. This multiallelic mating locus contains two genes (*b-East* and *b-West*), which code for a pair of homeodomain transcription factors. Upon mating of compatible cells, pathogenic and sexual development are triggered by a heterodimeric bE/bW complex (10). Since the *MbA* genome is completely equipped with mating type genes, we first deployed a screen for potential mating partners. To this end, we screened wild *M. bullatus* isolates to find a suitable mating partner, but we could not observe any mating event (Supplementary Figure S7). To test if *MbA* is able to undergo pathogenic differentiation in the absence of mating, we generated a self-compatible strain (CB1) which carries compatible b-mating alleles: to construct the CB1 strain, we used compatible alleles of the *b-East* and *b-West* genes of the barley smut *U. hordei*, a pathogen which is the phylogenetically most closely related to *MbA* and amenable to reverse genetics. The native *MbA* locus was replaced by the compatible *U. hordei b-East* and *b-West* gene alleles via homologous recombination (Figure 6B).

**Figure 6:**
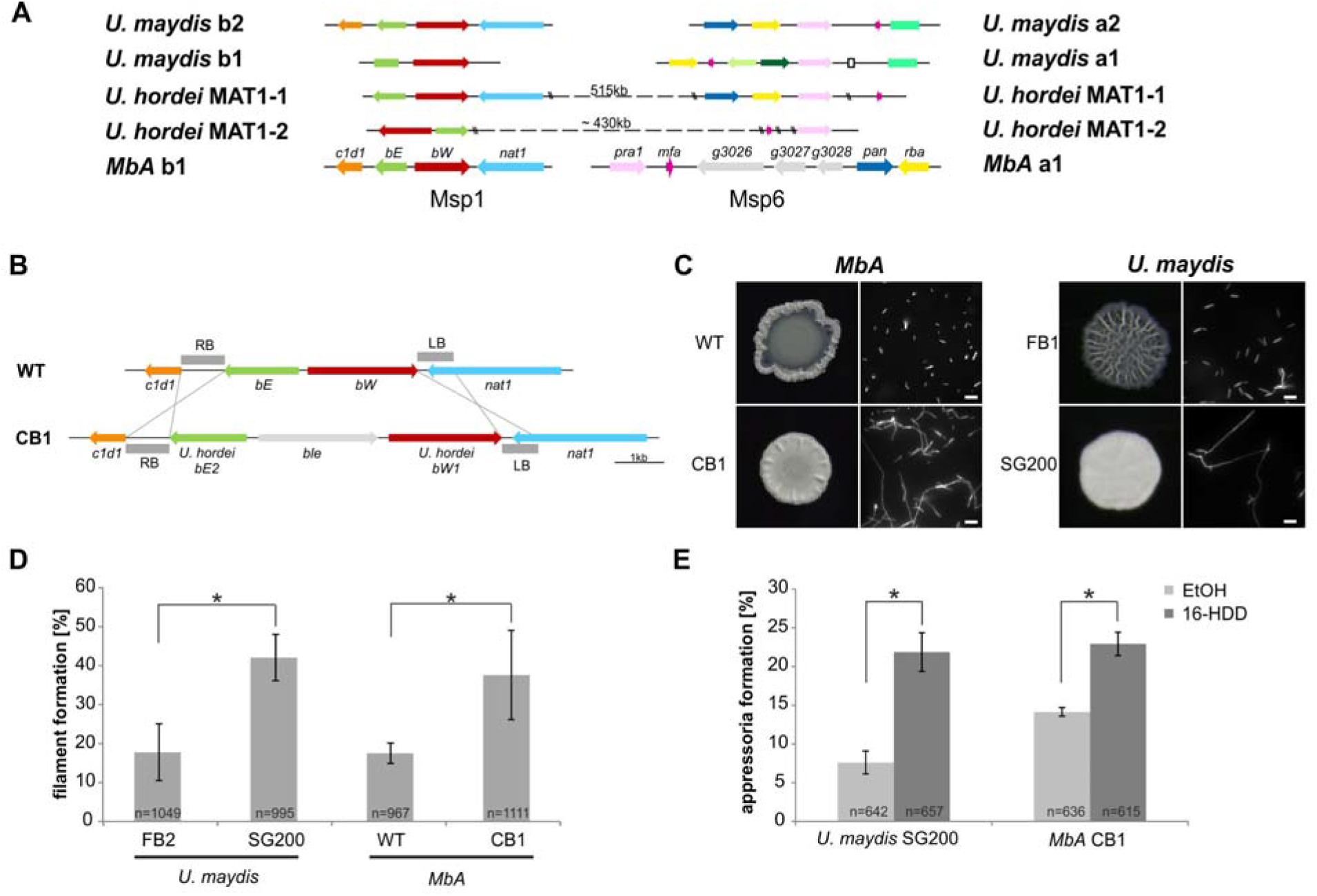
The self-compatible *MbA* strain CB1 (A) *MbA* mating type genes, unlike the ones of *U. hordei*, can be found on two different chromosomes similar to the tetrapolar mating type system of *U. maydis*. (B) To generate a self-compatible strain (CB1), the b-mating genes of *U.hordei* were integrated at the native *MbA b-*locus. (C) Unlike the *MbA* wild type strain (top left), strain CB1 (bottom left) shows a fluffy phenotype on charcoal plates and filamentous growth. *U. maydis* haploid F1 strain (top right) and self-compatible SG200 strain (bottom right) were used as negative and positive control, respectively. (D,E) Induction of filamentation and appressoria formation in strain CB1 was studied in three independent experiments. For this around 1000 cells for filament formation and around 600 cells for appressoria formation were analyzed and error bars indicate standard error. After incubation on a hydrophobic surface, both, filament and appressoria formation in strain CB1, were significantly different (* Chi-Square Test for Independence – α = 0,0001) when compared to *MbA* wild type and similar to the level of the self-compatible *U. maydis* strain SG00. *U. maydis* haploid F2 strain was used as negative control (18). Scale bar: 20μm. 16-HDD: 16-Hydroxyhexadecanoic acid.

Incubation of the *MbA* CB1 on charcoal plates led to the formation of aerial hyphae with the characteristic fluffy phenotype of filamentous strains like the self-compatible, solopathogenic *U. maydis* SG200 strain (Figure 6C). A second established method to induce filament formation in smuts is on hydrophobic parafilm (27). Quantification after hours incubation of *MbA* CB1 on parafilm resulted in the formation of filaments comparable to those of the *U. maydis* SG200 strain (Figure 6D). While about 17% of *MbA* wild type cells showed filaments, the CB1 strain with compatible b-genes showed 38% filamentous growth.

Formation of appressoria is a hallmark of pathogenic development in smut fungi (27). While the switch from yeast-like growth to filamentous development is the first step in the pathogenic development of smut fungi, host penetration is accompanied by the formation of a terminal swelling of infectious hyphae, termed “appressoria”. Induction of appressoria-formation *in vitro* can be induced by adding 100 μM of the cutin monomer 16-Hydroxyhexadecanoic acid (HDD) to the fungal cells prior to cell spraying onto a hydrophobic surface (27). In absence of HDD, only about 8% of the *U. maydis* SG200 cells and 14% of the *MbA* cells formed appressoria on parafilm 24 hours after spraying (Figure 6E). Addition of 100μM HDD resulted in a significant induction of appressoria in both *U. maydis* and *MbA*, demonstrating that *MbA* does hold the genetic repertoire to form infection structures *in-vitro*. Together, the analysis of the recombinant CB1 strain indicates that *MbA* can sense pathogenesis-related surface cues and produce penetration structures to a similar level as that seen for the pathogenic model organism *U. maydis*.

### Identification of microbe-microbe effector genes by RNA-Seq

To study the transcriptomic response of *MbA* to different biotic interactions, RNA sequencing was performed. The *MbA* transcriptome was profiled in five different conditions (Figure 7A; cells in axenic culture versus cells on-planta, on-planta + SynCom, on-planta + *A. laibachii*, on-planta + SynCom + *A. laibachii*). Inoculations of *A. thaliana* leaves were performed as described above for *A. laibachii* infection assays (Figure 7A). For *MbA* RNA preparation, the epiphytic microbes were peeled from the plant tissue by using liquid latex (see methods section for details).

**Figure 7:**
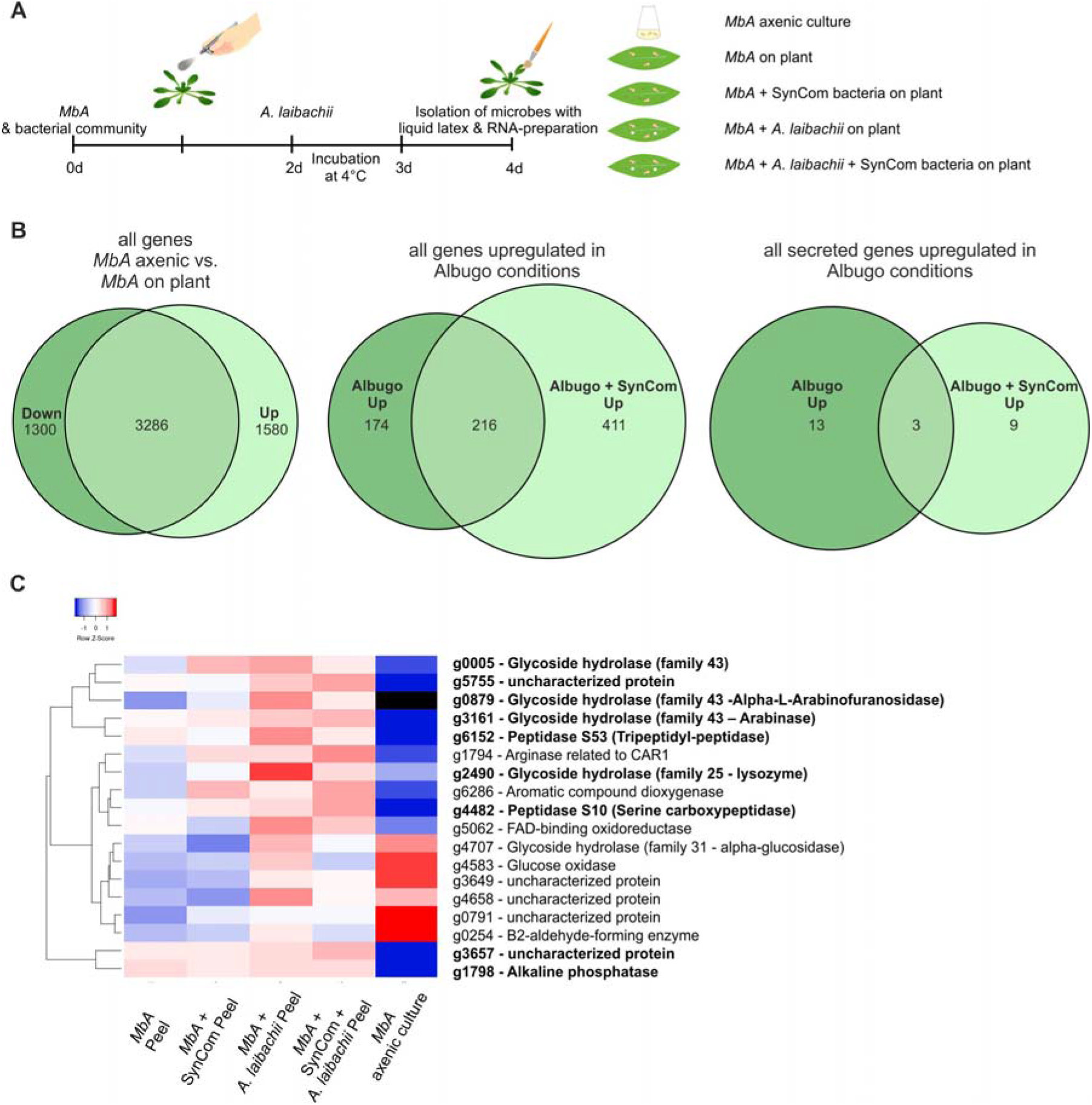
Transcriptome analysis of *MbA* (A) Experimental setup used for the transcriptomic (RNA-Sequencing) analysis in *MbA*. (B) Venn diagrams showing differential regulated *MbA* genes after spraying of haploid cells onto the *A. thaliana* leaf surface. A total number of 801 genes were upregulated in response to *A. laibachii* in presence and absence of bacterial SynCom. 216 of the 801 genes were upregulated in both conditions. (C) Hierarchical clustering of the 27 *A. laibachii* - induced *MbA* genes that are predicted to encode secreted proteins. Of these genes, nine were selected as candidate microbe-microbe effector genes, based on their transcriptional upregulation and prediction to encode for extracellularly localized proteins.

The libraries of the 15 samples (five conditions in three biological replicates each) were generated by using a poly-A enrichment and sequenced on an Illumina HiSeq4000 platform. The paired end reads were mapped to the *MbA* genome by using Tophat2 (28). The analysis revealed that *MbA* cells on *A. thaliana* leaves (on-planta) downregulated 1300 and upregulated 1580 genes compared to cells in axenic culture (Figure 7B). Differentially expressed genes were determined with the “limma”-package in R on “voom” (Supplementary Figure S8) using a False discovery rate threshold of 0.05 and log2FC > 0. A GO-terms analysis revealed that, among the downregulated genes, 50% were associated with primary metabolism (Supplementary Figure S9). In the two conditions in which *A. laibachii* was present, we observed upregulation of 801 genes. Among these genes, 411 genes were specific to co-incubation of *MbA* with *A. laibachii* and SynCom while 174 were specific to incubation with *A. laibachii* only. A set of 216 genes was shared in both conditions (Figure 7B).

In presence of *A. laibachii,* mainly metabolism- and translation-dependent genes were upregulated, which might indicate that *MbA* can access a new nutrient source in presence of *A. laibachii* (Supplementary Figure S9). Among all *A. laibachii* – induced *MbA* genes, 25 genes encode proteins carrying a secretion signal peptide and having no predicted transmembrane domain (Figure 7C). After excluding proteins being predicted to be located in intracellular organelles, nine candidate genes remained as potential microbe-microbe dependent effectors, i.e. *MbA* genes which are induced by *A. laibachii*, show no or low expression in axenic culture and encode for putative secreted proteins (Figure 7C). Interestingly, four of these genes encode putative glycoside hydrolases. Furthermore, two genes encode putative peptidases, one gene likely encodes an alkaline phosphatase and two encode uncharacterized proteins (Figure 7C).

To directly test the eventual antagonistic function of those genes towards *A. laibachii*, we selected the two predicted glycoside hydrolases-encoding genes *g5* & *g2490* (GH43 & GH25) and the gene encoding the uncharacterized protein *g5755* for gene deletion in *MbA*. The respective mutant strains were tested in stress assays to assess, whether the gene deletions resulted in general growth defects. Wild type and mutant *MbA* strains were exposed to different stress conditions including osmotic stress (sorbitol, NaCl), cell wall stress (calcofluor, congored) and oxidative stress (H_2_O_2_). Overall, in none of the tested conditions we observed a growth defect of the deletion mutants in comparison to wild type *MbA* (Supplementary Figure S10). To test an eventual impact of the deleted genes in the antagonism of the two microbes, the *MbA* deletion strains were each pre-inoculated on *A. thaliana* leaves prior to *A. laibachii* infection. Deletion of *g5* resulted in a significant but yet marginal increase of *A. laibachii* disease symptoms, while deletion of *g5755* had no effect on *A. laibachii*. We therefore considered these two genes being not important for the antagonism of *MbA* towards *A. laibachii.* Strikingly, the *MbA* Δ*g2490* strain almost completely lost its biocontrol activity towards *A. laibachii*. This phenotype was reproduced by two independents *g2490* deletion strains (Figure 8A). To check if this dramatic loss of microbial antagonism is specific to the deletion of *g2490*, in-locus genetic complementation of strain Δ*g2490_1* was performed via homologous recombination. The resulting strain *MbA* Δ*g2490/compl* regained the ability to suppress *A. laibachii* infection, confirming that the observed phenotype specifically resulted from the deletion of the *g2490* gene (Figure 8B). Together, these results demonstrate that the biocontrol of the pathogenic oomycete *A. laibachii* by the basidiomycete yeast *MbA* is determined by the secretion of a previously uncharacterized GH25 enzyme, which is transcriptionally activated specifically when both microbes are co-colonizing the *A. thaliana* leaf surface.

**Figure 8:**
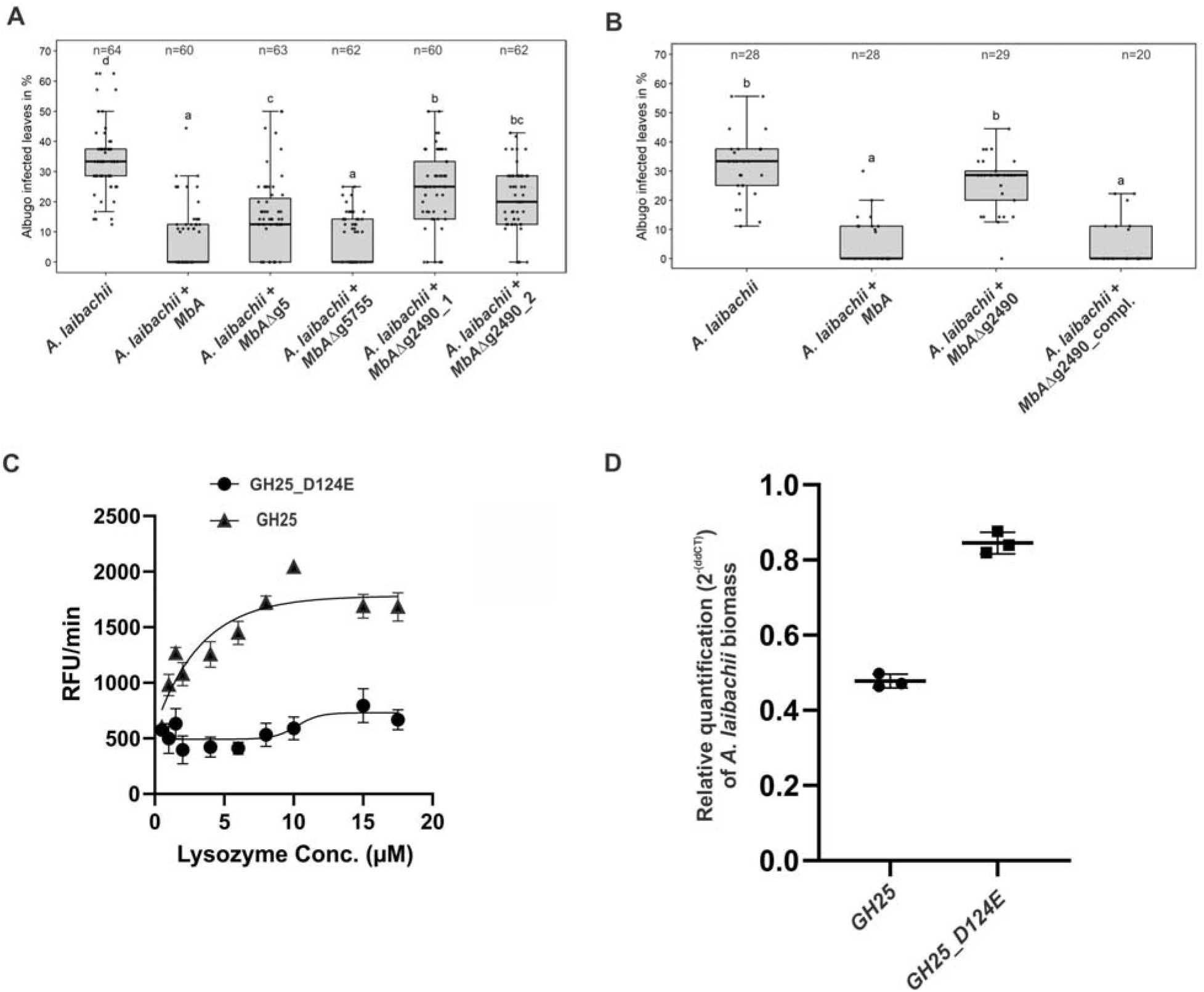
A reverse-genetic approach to identify the *MbA* gene which is responsible for the suppression of *A. laibachii* infection. (A) Three candidate microbe-microbe effector genes (*g5*, *g5755* &*g2490*) were deleted in *MbA* and deletion strains were individually inoculated on *A. thaliana* together with *A. laibachii*. Inoculation of two independent *g2490 null* strains (Δ*g2490_1;* Δ*g2490_2)* resulted in significant and almost complete loss the biocontrol activity of *MbA*. While deletion of g5 resulted in a marginal reduction of disease symptoms at 14 days post infection, deletion of g5755 had no effect on *A. laibachii*. (B) Genetic complementation of the *g2490* deletion restores the biocontrol activity to wild type levels. Infections in (A) were performed in six, in (B) in three individual replicates. In each replicate 12 plants were infected. N indicates the number of infected plants that were scored for symptoms. Different letters indicate significant differences (P values < 0.05; ANOVA model for pairwise comparison with Tukey’s HSD test). (C) Detection of lysozyme. Increasing concentrations of purified MbA_GH25 and MbA_GH25(D124E) were incubated with the DQ lysozyme substrate for an hour at 37 °C. The fluorescence was recorded every minute in a fluorescence microplate reader using excitation/emission of 485/530 nm. Finally, Relative Fluorescence Unit (RFU)/ min was calculated for each concentration and plotted on the graph. Each data point represents three technical replicates and three independent biological replicates as indicated by the Standard Error Measurement (SEM) bars. An unpaired t-test was performed for the active GH25 and Mutant_GH25 sets giving the p-value of <0.0001; and R squared value of 77.24%. (D) Relative quantification of *A. laibachii* biomass in response to MbA_GH25 (active and mutant) treatment via qPCR. The Oomycete internal transcribed spacer (ITS) 5.8s, was normalized to *A. thaliana* EF1-α gene to quantify the amount of *A. laibachii* DNA in the samples, ten days post infection. Then relative biomass was calculated comparing control sets (Only Albugo) with *A. laibachii* treated with GH25 and *A. laibachii* treated with Mutant_GH25 by ddCT method. Unpaired t-test between GH25 and Mutant_GH25 sets gave a p-value of <0.0001 and an R-squared value of 98.88%.

### Functional characterization of the secreted MbA hydrolase

To characterize the protein function of the GH25 encoded by *MbA g2490*, we were using *Pichia pastoris* for heterologous expression. The recombinant protein was tagged with polyhistidine tag for Ni-NTA affinity purification. The purified protein was detected at an expected size of 27kDa (Supplementary Figure S11A). In addition, via site directed mutagenesis a mutated version of the protein was generated, carrying a single amino exchange at the predicted active site (GH25_D124E). Both active and mutated versions of the GH25 hydrolase were subjected to a quantitative lysozyme activity assay using fluorogenic substrate *Micrococcus lysodeikticus* with commercial Hen egg-white lysozyme as a control. We noticed a concentration dependent increase in relative fluorescence unit (RFU)/min for the active GH25 in molar concentrations from 2uM to 10uM. Whereas, for similar concentrations, mutated GH25 (GH25mut) showed no significant increase in RFU/min compared to the active version. Commercial HEWL showed a steady increase in RFU/min from 1uM to 5.5uM concentrations (Figure 8C; Supplementary Figure S11C). Thus, the recombinant protein represents a functional GH25 hydrolase with a lysozyme activity.

To test for a direct function of the GH25 lysozyme, we treated *A. laibachii* -infected Arabidopsis plants with the recombinant protein. The impact of treatment of *A. laibachii* infection was quantified by quantitative PCR to determine the relative *A. laibachii* biomass on Arabidopsis in response to GH25. Strikingly, we observed a significant reduction of *A. laibachii* colonization in leaves treated with the active GH25 lysozyme, while the mutated enzyme GH25_D124E did not significantly influence infection (p-value of <0.0001 and an R-squared value of 98.88%) (Figure 8D). Overall, treatment with the GH25 lysozyme reduced the amount of *A. laibachii* to about 50%.

## Discussion

Healthy plants in natural habitats are extensively colonized by microbes, therefore it has been hypothesized that the immune system and the microbiota may instruct each other beyond the simple co-evolutionary arms race between plants and pathogens (29). Community members as individuals or in a community context have been reported to confer extended immune functions to their plant host. Root endophytic bacteria for example were found to protect *A. thaliana* and stabilize the microbial community by competing with filamentous eukaryotes (30). A large inhibitory interaction network was found in the leaf microbiome of *A. thaliana* and genome mining was used to identify over 1000 predicted natural product biosynthetic gene clusters (BGCs) (31). In addition, the bacterium *Brevibacillus* sp. leaf 182 isolate was found to inhibit half of the 200 strains isolated from *A. thaliana* phyllosphere. Further analysis revealed that *Brevibacillus* sp. leaf 182 produces a trans-acyltransferase polyketide synthase-derived antibiotic, macrobrevin along with other putative polyketide synthases (31).

In this study, we describe the role of the basidiomycete yeast *MbA,* which we previously co-isolated with the oomycete pathogen *A. laibachii* and now characterized as an antagonistic driver in the *A. thaliana* phyllosphere. *A. laibachii* inhibits *in-vitro* growth of seven members of a bacterial leaf SynCom and, most strikingly, strongly suppresses disease progression and reproduction of the pathogenic oomycete *A. laibachii* on *A. thaliana*. *MbA* is a member of the Ustilaginales, which had previously been classified into the group of pathogenic smut fungi of the *Moesziomyces bullatus* species (11). Our genome analysis identified the anamorphic yeasts *M. rugulosus, M. aphidis* and *M. antarcticus*, which had previously been classified as “*Pseudozyma spec*.”, as the closest relatives of *MbA*. Anamorphic Ustilaginales yeasts are long known and have been used for biotechnological applications and also biocontrol (32). Mannosylerythritol lipids produced by *M. antarcticus* are known to act as biosurfactants and are of great interest for pharmaceutical applications (33,34). Glycolipids like flocculosin produced by *A. flocculosa* or ustilagic acid characterized in the smut fungus *U. maydis*, have antifungal activity. Those compounds destabilize the membrane of different fungi and thus serve as biocontrol agents against powdery mildews or grey mold (35–37)

We identified 13 potential secondary metabolite gene clusters in *MbA,* including non-ribosomal peptide synthases and polyketide synthetase. Interaction among microbes within the same habitat is believed to have given rise to a variety of secondary metabolites (38,39). The presence of *Streptomyces rapamycinicus* was shown to activate an otherwise silent polyketide synthase gene cluster, *fgnA*, in *Aspergillus fumigatus*. The resultant compound proved to be a potent fungal metabolite that inhibited the germination of *S. rapamycinicus* spores (40). Therefore, secondary metabolite gene clusters and their corresponding products may confer a competitive advantage to fungi over the bacteria that reside in the same environment.

What is still under debate is the relation of anamorphic yeasts with the related pathogenic smuts. Many smut fungi, including the model species *U. maydis* are dimorphic organisms. In their saprophytic phase they grow as haploid non-pathogenic yeast cells. Only on appropriate host surfaces, haploid cells switch to filamentous growth and expression of pathogenicity-related genes is only activated upon mating in the filamentous dikaryon. A prime prerequisite for pathogenic development is therefore the ability of mating (41,42). Our genome analysis identified a tetrapolar mating system with a complete set of mating genes in *MbA*. Looking more closely on the phylogeny of different mating genes it appears that all sequenced *Moesziomyces* strains have the same pheromone receptor type (Supplementary Figure S12). Together with our unsuccessful mating assays, this suggests that all sequenced strains of this species have the same mating type and, therefore, are unable to mate. Mating type bias after spore germination was reported for *Ustilago bromivora,* which leads to a haplo-lethal allele linked to the MAT-2 locus (43). In this case, an intratetrad mating event rescues pathogenicity in nature as the second mating partner is not viable after spore germination. Together with the observation that anamorphic *Moesziomyces* yeasts are ubiquitous in nature, one could hypothesize that these fungi are highly competitive in their haploid form and antagonism might have led to the selection of one viable mating type. This eventually adapted to the epiphytic life style.

Transcriptome analysis showed that epiphytic growth of *MbA* on *A. thaliana* leads to massive transcriptional changes particularly in primary metabolism, which might reflect adaptation to the nutritional situation on the plant surface. Moreover, *MbA* showed specific transcriptional responses to a bacterial community, as well as to *A. laibachii* when being co-inoculated on plant leaves. Presence of *A. laibachii* resulted in the induction of primary metabolism and biosynthesis pathways, which might reflect enhanced growth of *MbA* in the presence of *A. laibachii*.

A set of *MbA* genes encoding secreted hydrolases was induced by *A. laibachii* and one of these genes which encodes a putative GH25 hydrolase with similarity to *Chalaropsis* type lysozymes appeared to be essential for the biocontrol of *A. laibachii*. Initially discovered in the fungi *Chalaropsis* sp., this group of proteins is largely present in bacteria as well as phages for example the germination specific muramidase from *Clostridium perfringens* S40 (44). The bacterial muramidase, cellosyl from *Streptomyces coelicolor* (45) also belongs to the *Chalaropsis*type of lysozyme. These proteins are proposed to cleave the β-1,4-glycosidic bond between N-acetylmuramic acid (NAM) and N-acetylglucosamine (NAG) in the bacterial peptidoglycan. Specifically, the β -1,4-N,6-O-diacetylmuramidase activity allows the *Chalaropsis* type lysozyme to degrade the cell wall of *Staphylococcus aureus*, in contrast to the commercially available Hen egg-white lysozyme (HEWL) (45). Despite differences in structure and molecular weight from HEWL, the GH25 of MbA has lysozyme activity against the gram positive bacterium *Micrococcus lysodeikticus* in a fluorogenic assay. This highlights the overall biochemical functionality of the recombinant glycoside hydrolase. The glycoside Hydrolase 25 family is predicted to have an active site motif DXE which is highly conserved across the fungal kingdom (Supplementary Figure S13). The structure of glycoside hydrolase family 25 from *Aspergillus fumigates* was characterized and the presence of N-terminal signal peptide was considered to indicate an extracellular secretion of the protein with possible antimicrobial properties (46). The role of the secreted hydrolase in the fungal kingdom is not completely explored yet. The presence of such hydrolases has in many cases been hypothesized to be associated with hyperparasitism of fungi parasitizing fungi (47) or oomycetes parasitizing oomycetes (48). Our results might therefore indicate a cross kingdom hyperparasitism event between a fungus and an oomycete. Previous work on microbial communities has indicated that negative interactions stabilize microbial communities. Hyperparasitism is such a negative interaction with a strong eco-evolutionary effect on pathogen-host interactions and therefore on community stability (49). *MbA* might therefore regulate *A. laibachii* infection and reduce disease severity. The qPCR evaluation of oomycete biomass strongly points towards the idea that *A. laibachii* is a direct target of antagonism for MBA. Since we observed reduced formation of *A. laibachii* in presence of *MbA*, we also tested if the GH25 lysozyme would suppress zoospore germination. However, we could not detect a significant reduction of *A. laibachii* zoosporangia germination upon treatment with active GH25 lysozyme (Supplementary figure S14), suggesting that the GH25 lysozyme interferes with *A. laibachii* at a later stage of infection. As *A. laibachii* has been shown to reduce microbial diversity (7), *MbA* might increase diversity through hyperparasitism of *A. laibachii*. At the same time this increased diversity might have caused the need for more secondary metabolites to evolve in the *MbA* genome to defend against niche competitors. Through its close association with *A. laibachii*, *MbA* could be a key regulator of the *A. thaliana* microbial diversity and therefore relevant for plant health beyond the regulation of *A. laibachii* infection.

In conclusion, the secreted hydrolase we identified as main factor of *A. laibachii* inhibition has great potential to act as antimicrobial agent. The isolated compound is not only valuable per se in an ecological context. It can further lay the grounds for exploring other microbial bioactive compounds that mediate inter-species and inter-kingdom crosstalk. A main goal of our future studies will be to understand on the mechanistic level, how the GH-25 suppresses *A. laibachii*, and at which developmental step the oomycete infection is blocked. Since the GH-25 enzyme is well conserved amongst Ustilaginales including pathogenic species, it will also be tempting to elucidate whether the species-specific antagonism identified here is broadly conserved among Ustilaginales fungi and oomycetes. We further will investigate potential responses by the host plant and how this impacts *A. laibachii* growth upon *MbA* colonization. Functional investigation of these interactions can provide meaningful insights as to why certain yeasts prefer to colonize specific environments. At the same time, it will be worth exploring how the basidiomycete yeasts influence the bacterial major colonizers of the phyllosphere.

## Material and Methods

### Strains and growth conditions

*MbA* wildtype strain was isolated from *A. laibachii* infected *A. thaliana* leaves [7]. Wild-type *MbA* (at 22 degrees) and *U. maydis* (at 28 degrees) strains were grown in liquid YEPSlight medium and maintained on Potato dextrose agar plates. King’s B medium was used for culturing Syn Com bacterial members at 22 degrees. All the strains were grown in a rotary shaker at 200rpm. All the recipes for medium and solutions can be found in Supplementary Table S2. Stress assays for fungi: wildtype and mutant strains of *MbA* grown to an optical density (600 nm) of 0.6-0.8 were centrifuged at 3500rpm for 10 minutes and suspended in sterile water to reach an OD of 1.0. Next, a dilution series from 10^0^ to 10-4 was prepared in sterile H_2_O. In the end, 5 μl of each dilution were spotted on CM plates supplemented with the indicated stress agents. The plates were incubated for 2 days at 22°C. Confrontation assays: at first, *MbA* and SynCom bacterial strains were grown to an O.D of 0.8-1. *MbA* cultures (10ul) were dropped in four quadrants of a Potato Dextrose Agar plate, previously spread with a bacterial culture. Plates were incubated for 2-4 days at 22°C.

### Transformation of *MbA* and plasmid construction for generation of knockout mutants

Fungal strains were grown in YEPSL at 22°C in a rotary shaker at 200rpm until an O.D. of 0.6 was reached and centrifuged for 15 mins at 3500rpm. The cells were washed in 20 ml of SCS (Table S2), and further centrifuged for 10 minutes at 3000rpm, before being treated with 3ml SCS solution with 20mg/ml of Glucanex (Lysing Enzyme from *Trichoderma harzianum*, # L1412, Sigma). After 20 minutes of incubation at room temperature, as cell wall lysis was occurred, cold SCS was added to the mix and protoplasts spun down for 10 minutes at 2400rpm. They were then washed twice with SCS and resuspended with 10 ml STC (SupplementaryTable S2) to be centrifuged at 2000rpm for 10 minutes. Finally, the pellet was dissolved in 500 μl STC, and stored in aliquots of 50 μl at −80°C. 5μg of plasmid DNA along with 15 μg Heparin was added to 50 μl protoplasts. After incubation on ice for 10 minutes, STC/40%PEG (500 μl) was added to it and mixed gently by pipetting up and down; this step was followed by another 15 minutes on ice. The transformation mix was added to 10 ml of molten regeneration (reg) agar and poured over a layer of already solidified reg agar containing appropriate antibiotic solution. For the bottom layer, we used 400 μg/ml Hygromycin/ 8 μg/ml Carboxin/ 300 μg/ml nourseothricin (NAT).

Plasmids were cloned using *Escherichia coli* DH5α cells (Invitrogen, Karlsruhe, Germany). Construction of deletion mutants was performed by homologous recombination; the 5’ and 3’ flanking regions of the target genes were amplified and ligated to an antibiotic resistance cassette (50). The ligated fragment was subsequently transformed into *MbA*. Homologous integration of the target gene was verified via PCR on the antibiotic resistant colonies. Oligonucleotide pairs for knockout generation and verification can be found in Supplementary Table S3. PCR amplification was done using Phusion© DNA polymerase (Thermo Scientific, Bonn, Germany), following the manufacturer’s instructions, with 100 ng of genomic DNA or cDNA as template. Nucleic acids were purified from 1% TAE agarose gels using Macherey-Nagel™ NucleoSpin™ Gel and PCR Clean-up Kit.

### Mating assay and generation of the self-compatible *MbA* strain CB1

Haploid strains of *MbA* were grown in liquid cultures, mixed and drops arranged on PD-plates with charcoal to induce filament formation. Plate with the haploid *U. maydis* strains FB1 and FB2 and the solopathogenic strain SG200 served as internal control.

The complete b-locus of the solopathogenic *U. hordei* strain DS200 was amplified (Figure S2) and inserted into the *MbA* b-locus by homologous recombination. The strain obtained, known as compatible b1 (CB1) was tested positive by amplification of the right border and left border areas with primers specific for the genomic locus and for the plasmid region. Additionally, two primers specific for the *MbA bE*and *bW* genes were chosen to amplify parts of the native locus. To induce filament and appressoria formation in vitro we used a *Moesziomyces* YEPSL culture at OD_600_ 0.6-0.8. The cells were diluted to an OD_600_ of 0.2 in 2% YEPSL (for appressoria formation 100μM 16-hydroxyhexadecanoic acid (Sigma-Aldrich) or 1% ethanol was added) and sprayed the yeast like cells on parafilm which mimics the hydrophobic plant surface. After 18h incubation at 100% humidity the number of cells grown as filaments (or generating appressoria) was determined relative to the total number of total cells by using a light microscope.

### *Arabidopsis thaliana* leaf infections and quantification of Albugo biomass quantification by qPCR

Sterilized *Arabidopsis thaliana* seeds were subjected to cold treatment for 7 days and sown on 1/2 strength Murashige Skoog (MS) medium (Supplementary Table S2). The MS plates are directly transferred to growth chambers having 22°C on a short-day period (8 h light) with (33-40%) humidity and grown for 4 weeks before inoculation. Overnight liquid cultures of *MbA* and SynCom bacterial strains were grown to an OD_600_ of 0.6. The cultures were spun down at 3500rpm for 10 minutes and the pellets dissolved in MgCl_2_. 500μl of each culture was evenly sprayed on three-week old *A. thaliana* seedlings using airbrush guns. Two days later, a spore solution of *A. laibachii* was then sprayed on the seedlings following the protocol of Ruhe et al. (51). Two weeks later, the disease symptoms on the leaves were scored as a percentage between infected and non-infected leaves.

4 weeks old *A. thaliana* seedlings on MS plates were sprayed with *A. laibachii* as a control and GH25+ *A.laibachii* and *Mut_GH25+A.laibachii* as treatments. After 10 days post infection (dpi), the seedlings were harvested, frozen in liquid nitrogen and kept at −80°C. For DNA extraction, the frozen plant material was ground into a fine powder with mortar and pestle and treated with extraction buffer (50 mM Tris pH 8.0, 200 mM NaCl,0.2 mM ethylenediaminetetraacetic acid (EDTA), 0.5% SDS,0.1 mg/ml proteinase K (Sigma–Aldrich). This was followed centrifugation after the addition of one volume Phenol/Chloroform/Isoamylalkohol 25:24:1 (Roth). The top aqueous layer was removed and added to one volume of Isopropanol to precipitate the nucleic acids. DNA pellet obtained after centrifugation was washed with 70% EtOH and finally dissolved in 50 ul Nuclease-free water. For qPCR measurements; 10 ul of GoTaq® qPCR 2x Master Mix was used (Promega, Waltham, Madison, USA); 5ul of DNA (~50ng); 1ul of forward and reverse primer (10μM) up to a total volume 20μl. Samples were measured in triplicates in a CFX Connect real-time PCR detection system (Bio-Rad) following protocol of Ruhe et al. (2016)(51). Amount of *A. laibachii* DNA was quantified using the following oligonucleotide sequences, (A. thalianaEF1-α: 5′-AAGGAGGCTGCTGAGATGAA-3′, 5′-TGGTGGTCTCGAACTTCCAG-3′; Oomycete internal transcribed spacer (ITS) 5.8s: 5′-ACTTTCAGCAGTGGATGTCTA-3′, 5′-GATGACTCACTGAATTCTGCA-3′). Cq values obtained in case of the oomycete DNA amplification was normalized to *A. thalaina* DNA amplicon and then the difference between control (only *Albugo*) and treatment (*Albugo*+ GH25/Mut_GH25) was calculated by ddCq. The relative biomass of *Albugo* was analyzed by the formula (2^−ddCq^). Each data point in the graph represent three independent biological replicates.

### Nucleic acid methods

RNA-Extraction of Latex-peeled samples: Four weeks old *A. thaliana* plants were fixed between two fingers and liquid latex was applied to the leaf surface by using a small brush. The latex was dried using the cold air option of a hair dryer, carefully peeled off with a thin tweezer and immediately frozen in liquid nitrogen. Afterwards, the frozen latex pieces were grinded with liquid nitrogen and the RNA was isolated by using Trizol^®^ Reagent (Invitrogen, Karlsruhe, Germany) according to the manufacturer’s instructions. Turbo DNA-Free™ Kit (Ambion, life technologies^TM^, Carlsbad, California, USA) was used to remove any DNA contamination in the extracted RNA. Synthesis of cDNA was performed using First Strand cDNA Synthesis Kit (Thermo Fischer scientific, Waltham, Massachusetts, USA) according to recommended instruction starting with a concentration of 10μg RNA. QIAprep Mini Plasmid Prep Kit (QIAGEN, Venlo, Netherlands) was used for isolation of plasmid DNA from bacteria after the principle of alkaline lysis. Genomic DNA was isolated using phenol-chloroform extraction protocol (18).

RT-qPCR oligonucleotide pairs were designed with Primer3 Plus. The oligonucleotide pairs were at first tested for efficiency using a dilution series of genomic DNA. The reaction was performed in a Bio-Rad iCycler system using the following conditions: 2 min at 95 °C, followed by 45 cycles of 30 s at 95 °C, 30 s at 61 °C and generation of melting curve between 65°C to 95°C.

### Bioinformatics and computational data analysis

Sequence assembly of *MbA* strains was performed using the HGAP pipeline (Pacific Biosciences). *MbA* genome was annotated with the Augustus software tool. Secretome was investigated using SignalP4.0. Analysis of functional domains in the secreted proteins was done by Inter-Pro Scan. AntiSmash was used to predict potential secondary metabolite clusters. RNA sequencing was done at the CCG-Cologne Center for Genomics by using a poly-A enrichment on an Illumina HiSeq4000 platform. The achieved paired end reads were mapped to the *MbA* and *A. thaliana* TAIR10 genome by using Tophat2 (28). RNA-Seq reads of *MbA* axenic cultures were used to generate exon and intron hints and to start a second annotation with Augustus. Heat-maps were performed using the heatmap.2 function of the package gplots (version 3.0.1) in r-studio (R version 3.5.1). An analysis of variance (ANOVA) model was used for pairwise comparison of the conditions, with Tukey&apos;s HSD test to determine significant differences among them (*P* values <0.05).

### Heterologous protein production and GH25 activity assay

The *Pichia pastoris* KM71H-OCH gene expression system was used to produce MBA_GH25 domain tagged with an N-terminal Pilyhistidine tah (6XHis) and a C-terminal peptide containing the c-myc epitope and a 6xHis tag. The His-MspGH25 cloned into pGAPZαA vector (Invitrogen, Carlsbad, CA, USA) under the control of a constitutive promotor with an α-factor signal peptide for secretion. Expression and purification of recombinant proteins were performed according to manufacturer’s instructions (Invitrogen Corporation, Catalog no. K1710-01): YPD medium supplemented with 100 μg ml^−1^ zeocin was used for initial growth of *P. pastoris* strains at 28°C and 200 rpm (for liquid cultures). Production of the recombinant protein was performed in 1 L buffered (100 mM Potassium phosphate buffer, pH 6.0) YPD medium with 2% Sucrose at 28°C for 24 hours with 200 rpm shaking. Next the protein was subjected to affinity purification with a Ni-NTA-matrix, according to manufacturer’s instructions (Ni-Sepharose™ 6 Fast-Flow, GE-Healthcare; Freiburg, Germany). After purification, the His-MspGH25 protein was dialyzed in an exchange buffer (0.1M NaPi, 0.1M Nacl, pH=7.5). The purified protein was kept in 100 μl aliquots at 4°C.

Site directed mutagenesis was performed on pGAPZα-His-MspGH25 vector according to the instructions of the QuikChange Multi Site-Directed Mutagenesis Kit (Agilent Technologies, Santa Clara, United States) with primers targeting nucleotides of the active site of GH25.

Purified Glycoside Hydrolase of *MBA* from P. pastoris was quantified according to a sensitive fluorescence-based method using Molecular Probes™ EnzChek™ Lysozym-Assay-Kit (ThermoFisher Scientific, Katalognummer: E22013). DQ lysozyme substrate (*Micrococcus lysodeikticus*) stock suspension (1.0mg/ml) and 1000units/ml Hen egg White Lysozyme (HEWL) stock solution were prepared according to the manufacturer. Molar concentration of the HEWL stock solution was calculated using the following website (https://www.bioline.com/media/calculator/01_04.html) and was found to be 11μM. Protein concentration of MspGH25 (both active and mutated version was measured in the Nanodrop 2000c spectrophotometer (Thermo Fischer scientific, Waltham, Massachusetts, USA) according to manufacturer’s instructions using 100 μl of sample after using 100 μl of the appropriate buffer as a blank control in glass cuvette. The molar concentrations of recombinant proteins were also calculated as above.

Starting the reaction 50μl of the DQ lysozyme substrate working suspension was added to each microplate well containing reaction buffer with either HEWL (in molar concentrations ranging from 0.1-5.5μM) or MspGH25 (in molar concentration from 0.5-17.5μM). Fluorescence intensity of each reaction was measured every 5min to follow the kinetic of the reaction at 37°C for 60min, using fluorescence microplate reader with fluorescein filter Tecan Infinite 200 Pro plate reader (Tecan Group Ltd., Männendorf, Switzerland). Digestion products from the DQ lysozyme substrate have an absorption maximum at ~494nm and a fluorescence emission maximum at ~518nm.

## Data availability

Genome information and RNA sequencing have been submitted to NCBI Genbank and are available under the following links: https://www.ncbi.nlm.nih.gov/geo/query/acc.cgi?acc=GSE148670

## Acknowledgments

This work was funded through by the Deutsche Forschungsgemeinschaft (DFG, German Research Foundation) under Germany’s Excellence Strategy EXC-2048/1, Project ID 390686111, and the DFG priority program SPP2125 “DECRyPT”. We are grateful to Marco Thines for generously providing *M. bullatus* wild type strains. We thank Libera Lo Presti for critically reading the manuscript and helpful comments and suggestion.

## Supporting information captions

**Figure S1**: Biocontrol activity of *MbA,* but not *U. maydis,* against bacterial SynCom members (7). Inhibition by *Moesziomyces* can be seen as a characteristic halo after 48hrs of co-incubation.

**Figure S2:** Trypan blue staining of *A. thaliana* leaves 15 days post infection with *A. laibachii*. (i) & (ii)- control set with only *A.laibachii* (error bar 50μm); (i)- Thick hyphal growth of *A.laibachii* on the leaf surface, (ii) appressoria formation can be seen (red arrow). (iii) & (iv) Treatment set (*MbA* sprayed two days before *Albugo* (iii): Zoospores aggregated together, with few of them forming hyphae (green box); in addition, short, broken hyphae visualized in some regions (red box), which have not been found in Control sets; (error bar- 200μm) (iv): A closer look at the broken hyphae, yellow arrow indicates germinating cysts of *A. laibachii* (error bar - 50μm)

**Figure S3**: Genome comparison of *MbA* and *Moesziomyces antarctica* T-34. Highlighted regions show that contigs with chromosomal rearrangements in MBA can be also found in the genome of the related species *Moesziomyces antarctica* T-34.

**Figure S4**: (A) Predicted secondary metabolite clusters in the genome of *MbA*. Most clusters have unpredictable functions, three belong to the type of terpene or nonribosomal peptide synthetase types and one is a polyketide synthetase cluster type I. (B) The gene cluster encoding for production of ustilagic acid,a well-studied secondary metabolite of smut fungi (37), is not present in the genome of *MbA*. (C) Out of the 13 predicted secondary metabolite clusters, three are unique to *MbA*. Cluster 2 is predicted to encode a terpene, cluster 8 is a cluster of unknown function and cluster 10 is predicted as NRPS cluster. Core biosynthetic genes are highlighted in red, additional biosynthetic genes in yellow and transport-related genes in blue, based on AntiSMASH predictions.

**Figure S5**: Protein alignment of the core effector Pep1 (52) from different Ustilaginomycetes. (A) Pep1 regions important for functionality are present in all the aligned sequences. Deletion of the *pep1* gene (UMAG_01987) in *U. maydis* leads to complete loss of virulence, which can be restored by complementing the deletion mutant with the *MbA pep1* gene (Ma1682). Infection of maize leaves was done as described in (19). Disease symptoms were scored at 12 days post infection in three independent biological replicates. n= number of plants infected

**Figure S6**: Comparison of known virulence clusters (18) between *U. maydis* and *MbA*. Numbers are gene numbers (*UMAG_NUMBER* for *U. maydis*; *gNUMBER* for *MbA)*.

**Figure S7:** Mating assays of *MbA* and different *M. bullatus* isolates. Mixing of haploid *U. maydis* strains FB1 & FB2 and the solopathogenic strain SG200 served as a positive control for mating on charcoal plates, wherein filamentous growth is indicated by white, fluffy appearance of colonies. Haploid wild type strains without mating partner don’t show a fluffy phenotype. For all combinations of *Moesziomyces* strains, no mating event resulting in filamentous growth on charcoal plates was observed.

**Figure S8**: A) Multi-dimensional scaling plot (MDS) plot based on the interactions of *Moesziomyces* sp. (M.sp.) / *MbA* in response to the SynCom bacteria and *Albugo laibachii* in three biological replicates. The MDS plot shows *Albugo* and non-*Albugo* samples grouping together based on gene-level logCPM. (B) Voom mean-variance trend of the dataset where points represent genes, and (C) sample-specific weights obtained from the *limma-voom* function. Colours represent three replicates for each treatment. Light blue: *MbA* on plant; Dark blue: *MbA* on plant + SynCom; Light green: *MbA* on plant + Albugo; Dark green: *MbA* on plant + Albugo + SynCom.

**Figure S9**: (A) Sequence distribution of Gene Ontology terms. 60% of all the genes that are downregulated in *MbA* on plant compared to axenic culture growth can be assigned to GO-terms related to metabolism and cell cycle. (B) In contrast, presence of *A. laibachii* leads to transcriptional activation of metabolic processes. 52% of all GO-terms associated with genes upregulated in presence of *A. laibachii* are related to metabolic processes.

**Figure S10**: Stress assay of *MbA* wild type and knockout mutants of gene (g5, g5755 and g2490) respectively, on CM medium and 2% Glucose (A) with different conditions (B: 100 μg/ml Calcofluor; C: 150 μg/ml Calcoflour; D: 1 mMH2O2; E:45 μg/ml Congo-red; F: 1 M NaCl; G: 1 M Sorbitol). The strains were dropped on the CM plates containing different stress supplements in a dilution series from 10^0^ to 10^−4^

**Figure S11:** A) Recombinant MbA_GH25 was produced and purified using the *Pichia pastoris* protein expression system. The purified protein was loaded in a 12% SDS gel for visualization of an expected molecular weight of 27kDa for His-Tagged GH25. B) Schematic diagram of the recombinant construct, where the GH25 domain from MBA is tagged with an N-terminal polyhistidine Tag and a C-terminal peptide containing the c-myc epitope and a polyhistidine tag. C) Detection of lysozyme activity for Commercial Hen-egg white lysozyme (stock solution, 11μM) using the EnzChek^1^ Lysozyme Assay Kit. The fluorescence was recorded every minute in a fluorescence microplate reader using excitation/emission of 485/530 nm in increasing concentrations from 0.1μM to 5.5 μM. Finally, Relative Fluorescence Unit (RFU)/ min was calculated for each concentration and plotted on the graph.

**Figure S12:** A molecular phylogenetic analysis using maximum likelihood estimation and based on pheromone receptor protein sequences similarity. *MbA* protein sequence clusters together with type 1 pheromone receptors of other Ustilaginomycetes.

**Figure S13:** Amino acid alignment of GH25 sequences from different fungi (see attached list –‘GH25 with accession number’, for full length sequences). The protein sequences were obtained from the NCBI database. Alignment was achieved using the PRALINE multiple sequence alignment program with default parameters. The scoring scheme works from 0 for the least conserved alignment position, up to 10 (indicated by *) for the most conserved alignment position. A conserved active-site DxE motif has been predicted for glycoside hydrolase family 25. Sequences tested from different basiodiomycete, ascomycete and Chytrids, have the active site residue conserved (purple box).

**Figure S14:** Boxplot-analysis of GH 25 treatment on *in vitro A. laibachii* zoosporangial germination in three biological replicates analyzing about 100 zoosporangial cells for each replicate. A p value of 0.3 was obtained for paired T-test using one-tailed distribution.

**Table S1:** Composition of the bacterial SynCom

**Table S2**: *MbA* gene expression data

**Table S3**: Growth media and buffers used in this study

**Table S4**: PCR primers used in this study

