## Supplementary Figures for "A fungal member of the *Arabidopsis thaliana* phyllosphere antagonizes *Albugo laibachii* via a secreted lysozyme"

**Figure S1**

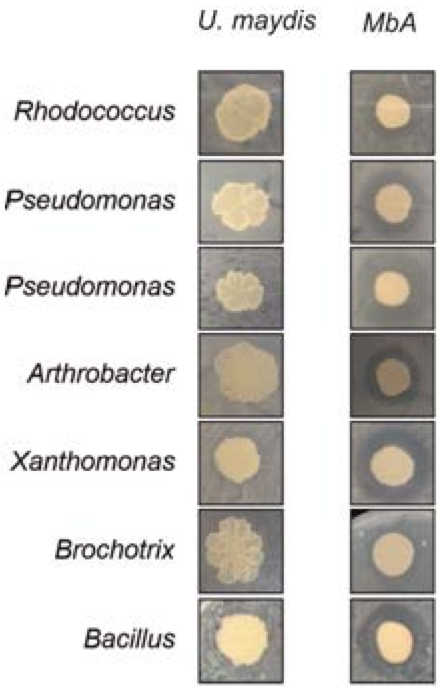


**Figure S1**: Biocontrol activity of *MbA* but not *U. maydis* against bacterial SynCom members [1]. Inhibiton by *Moesziomyces* can be seen by a characteristic halo formation after 48hrs of co-incubation.

**Figure S2**

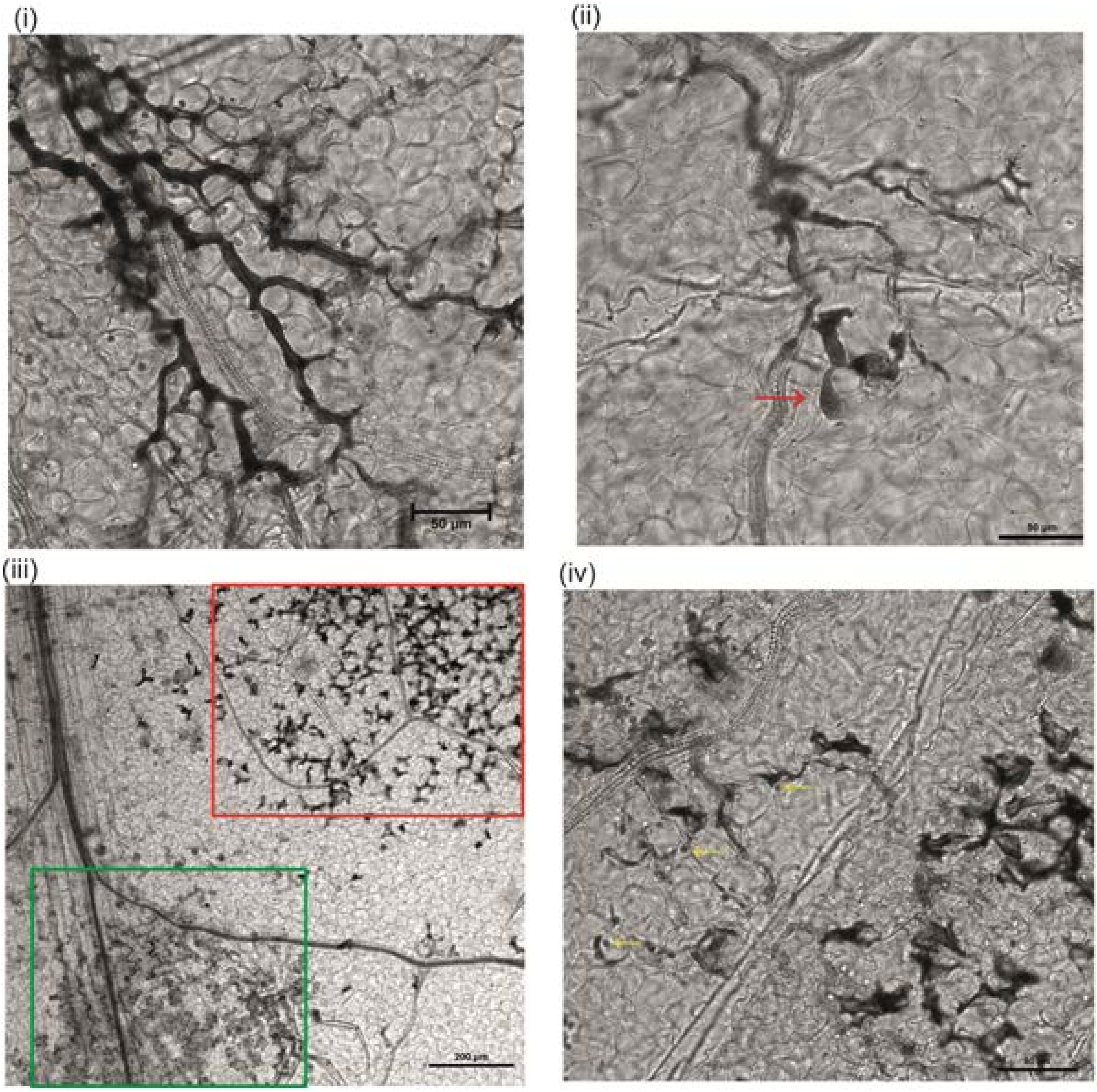


**Figure S2:** Trypan blue staining of *A. thaliana* leaves 15 days post infection with *A. laibachii*. (i) & (ii)- control set with only *A.laibachii* (error bar 50µm); (i)- Thick hyphal growth of *A. laibachii* on the leaf surface, (ii) appressoria formation can be seen (red arrow). (iii) & (iv) Treatment set (*MbA* sprayed two days before *A. laibachii* (iii): Zoospores aggregated together, with few of them forming hyphae (green box); in addition, short, broken hyphae visualized in some regions (red box), which have not been found in Control sets; (error bar-200µm) (iv): A closer look at the broken hyphae, yellow arrow indicates germinating cysts of *A. laibachii* (error bar - 50µm)

### Figure S3:

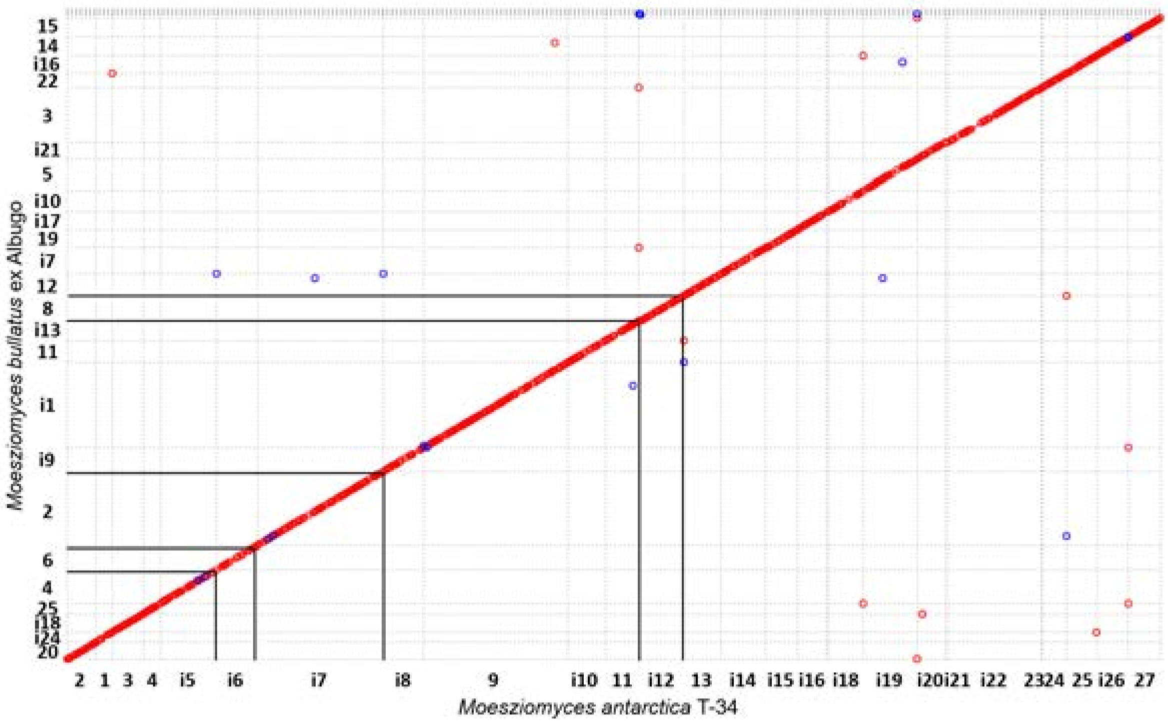


**Figure S3**: Genome comparison of MbA and *Moesziomyces antarctica* T-34. Highlighted regions show that contigs with chromosomal rearrangements in MBA can be also found in the genome of the related species *Moesziomyces antarctica* T-34.

**Figure S4**

A

| Cluster | Contig | Gene cluster type | Present in U. maydis |
|---|---|---|---|
| 1 | 1 | other | ✓ |
| 2 | 2 | Terpene cluster | X |
| 3 | 3 | other | ✓ |
| 4 | 3 | other | ✓ |
| 5 | 4 | other | ✓ |
| 6 | 5 | other | ✓ |
| 7 | 13 | Terpene cluster | ✓ |
| 8 | 17 | other | X |
| 9 | 20 | Nonribosomal peptide synthetase cluster | ✓ |
| 10 | 21 | Nonribosomal peptide synthetase cluster | X |
| 11 | 22 | Nonribosomal peptide synthetase cluster | ✓ |
| 12 | 24 | Type I PKS cluster | ✓ |
| 13 | 25 | Terpene cluster | ✓ |

B

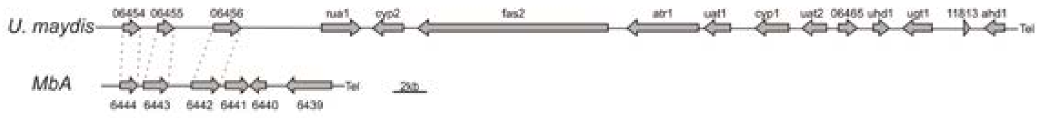


C

Cluster 2 - Terpene cluster

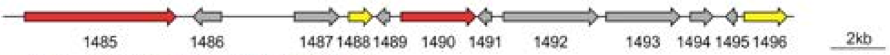


| gene | predicted function based on Blast2Go Annotation |
|---|---|
| 1485 | related to ser/thr protein kinase |
| 1486 | - |
| 1487 | related to translation initiation protein SUA5 |
| 1488 | probable theta class glutathione s-transferase |
| 1489 | probable Protein kish-A |
| 1490 | probable lanosterol synthase |
| 1491 | - |
| 1492 | related to RMD8 - Cytosolic protein required for sporulation |
| 1493 | tRNA-dihydrouridine(47) synthase [NAD(P)(+)] |
| 1494 | probable PPA1 - H+-ATPase 23 KD subunit, vacuolar |
| 1495 | related to 50S ribosomal protein L34 |
| 1496 | probable PRO2 - gamma-glutamyl phosphate reductase |

Cluster 8 - unknown cluster

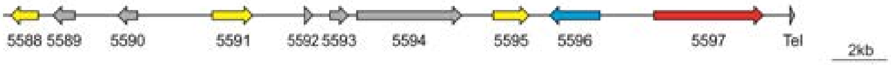


| gene | predicted function based on Blast2Go Annotation |
|---|---|
| [illegible] | [illegible] |
| 5589 | - |
| 5590 | Glutathione-dependent formaldehyde-activating enzyme |
| 5591 | Short-chain dehydrogenases/reductases |
| 5592 | - |
| 5593 | - |
| 5594 | fungal transcription factor regulatory middle homology region |
| 5595 | Aldo-keto reductases |
| 5596 | Sugar phosphate permease |
| 5597 | Thioester reductase domain of alpha aminoadipate reductase Lys2 and NRPSs Acyl-CoA synthetase (AMP-forming)/AMP-acid ligase II Phosphopantetheine attachment site |

Cluster 10 - NRPS cluster

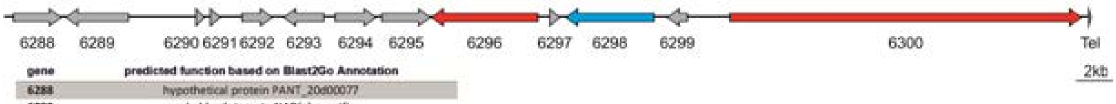


| gene | predicted function based on Blast2Go Annotation |
|---|---|
| 6288 | hypothetical protein PANT_20d00077 |
| 6289 | probable glutamate NAD(+)-specific |
| 6290 | 60S ribosomal L17 [Moesziomyces antarcticus] |
| 6291 | 60S ribosomal L17 [Moesziomyces antarcticus] |
| 6292 | probable RPL17A-ribosomal |
| 6293 | probable PAP1-poly(A) polymerase |
| 6294 | proteasome subunit alpha type 5 [Moesziomyces antarcticus] |
| 6295 | probable PUP2-20S proteasome subunit (alpha5) |
| 6296 | non-ribosomal peptide synthetase |
| 6297 | FOG: Zn-finger |
| 6298 | P-loop containing nucleoside triphosphate hydrolase |
| 6299 | related to C2H2 transcriptional regulator |
| 6300 | acetyl- synthetase |

**Figure S**4: (A) Predicted secondary metabolite clusters in the genome of *MbA*. Most clusters are having unpredictable functions, three are each belonging to the type of terpene or non-ribosomal peptide synthetase types and one is a polyketide synthetase cluster type I. (B) The gene cluster encoding for production of ustilagic acid, a well-studied secondary metabolite of smut fungi [2], is not present in the genome of *MbA*. (C) Out of the 13 predicted secondary metabolite clusters clusters, three are unique to *MbA*. Cluster 2 is predicted to encode a terpene, cluster 8 is a cluster of unkown function and cluster 10 is predicted as NRPS cluster. Core biosynthetic genes are highlighted in red, additional biosynthetic genes in yellow and transport related genes in blue, based on AntiSMASH predictions.

### Figure S5

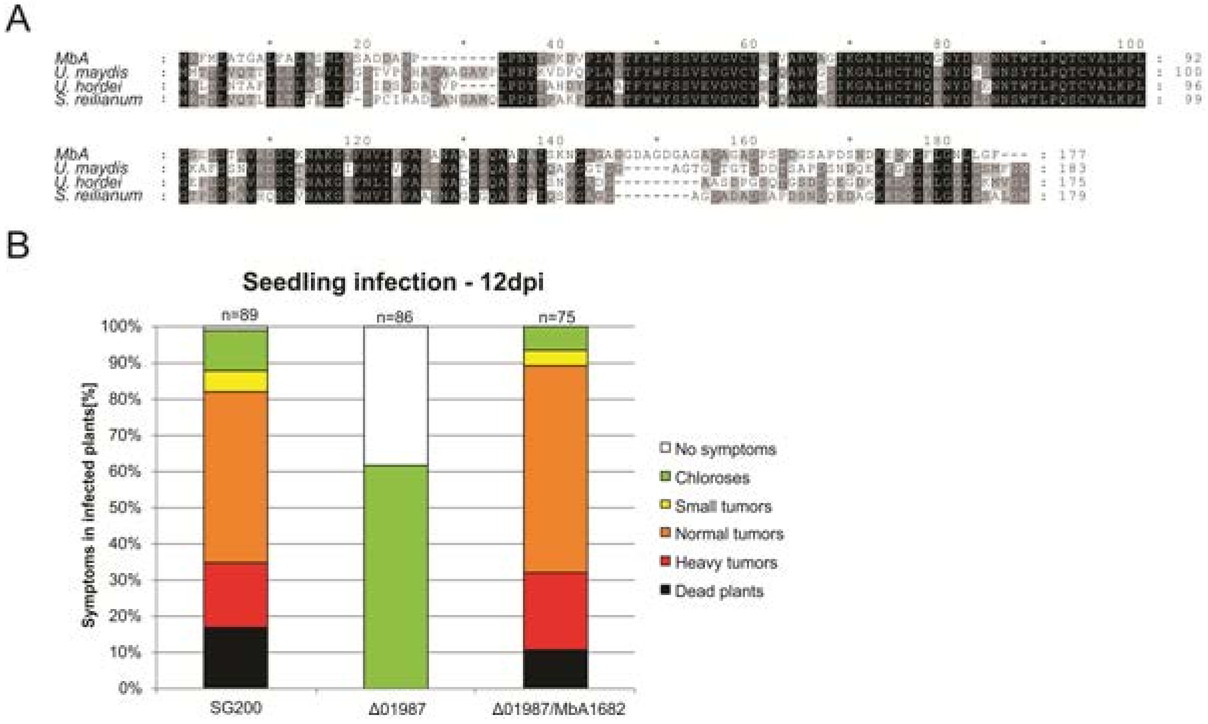


**Figure S5**: Protein alignment of the core effector Pep1 (Hemetsberger et al, 2015) from different Ustilaginomycetes (A). All characterized regions important for functionality are present in all different sequences. Deletion of the *pep1* gene (UMAG_01987) in *U. maydis* leads to complete loss of virulence, which can be restored by complementing the deletion mutant with the *MbA pep1* gene (Ma1682). Infection of maize leaves was done as described in [3]. Disease symptoms were scored at 12 days post infection in three independent biological replicates. n= number of plants infected

**Figure S6**

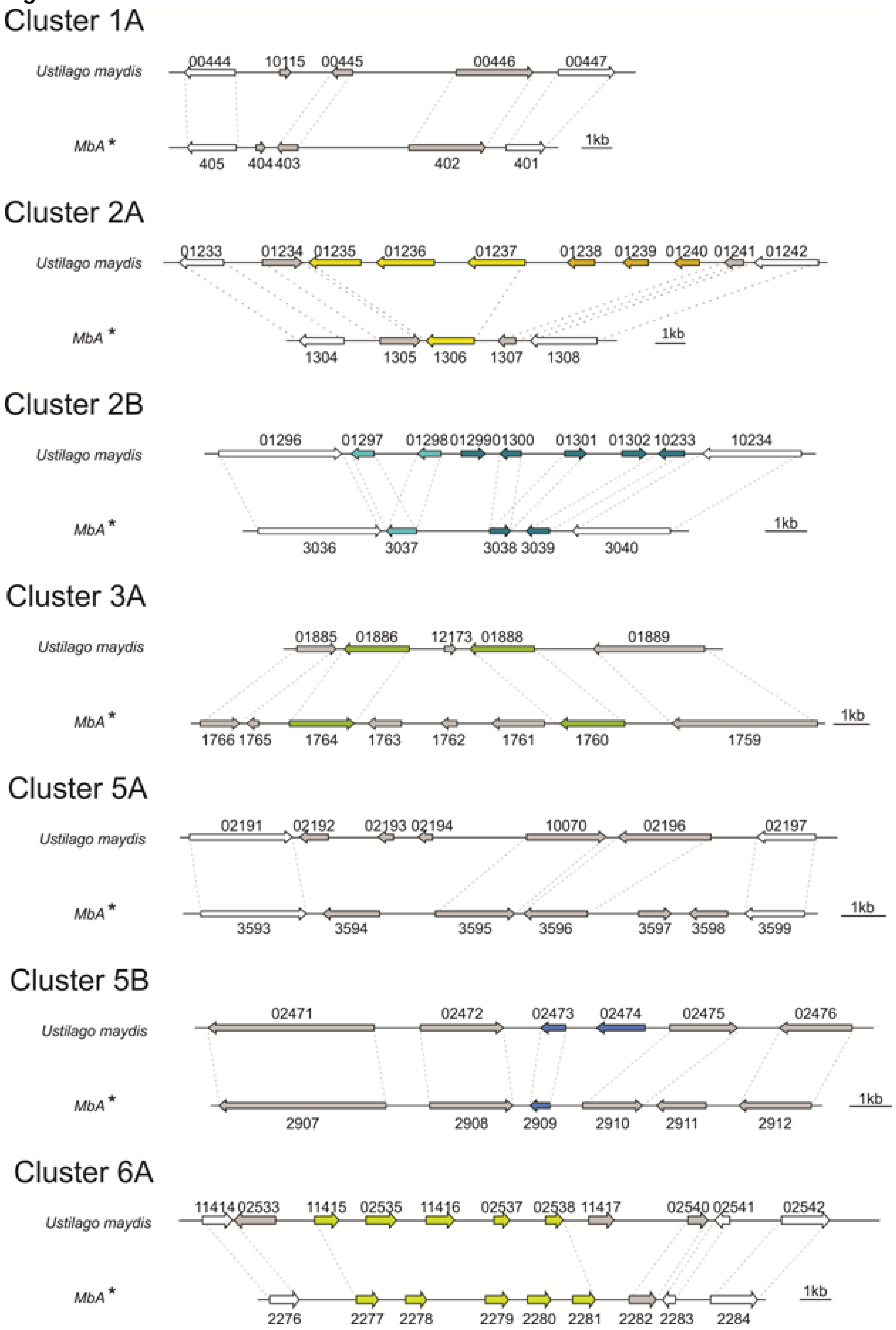
Cluster 1A
Ustilago maydis
00444 10115 00445 00446 00447
MbA *
405 404 403 402 401
1kb
Cluster 2A
Ustilago maydis
01233 01234 01235 01236 01237 01238 01239 01240 01241 01242
MbA *
1304 1305 1306 1307 1308
1kb
Cluster 2B
Ustilago maydis
01296 01297 01298 01299 01300 01301 01302 10233 10234
MbA *
3036 3037 3038 3039 3040
1kb
Cluster 3A
Ustilago maydis
01885 01886 12173 01888 01889
MbA *
1766 1765 1764 1763 1762 1761 1760 1759
1kb
Cluster 5A
Ustilago maydis
02191 02192 02193 02194 10070 02196 02197
MbA *
3593 3594 3595 3596 3597 3598 3599
1kb
Cluster 5B
Ustilago maydis
02471 02472 02473 02474 02475 02476
MbA *
2907 2908 2909 2910 2911 2912
1kb
Cluster 6A
Ustilago maydis
11414 02533 11415 02535 11416 02537 02538 11417 02540 02541 02542
MbA *
2276 2277 2278 2279 2280 2281 2282 2283 2284
1kb

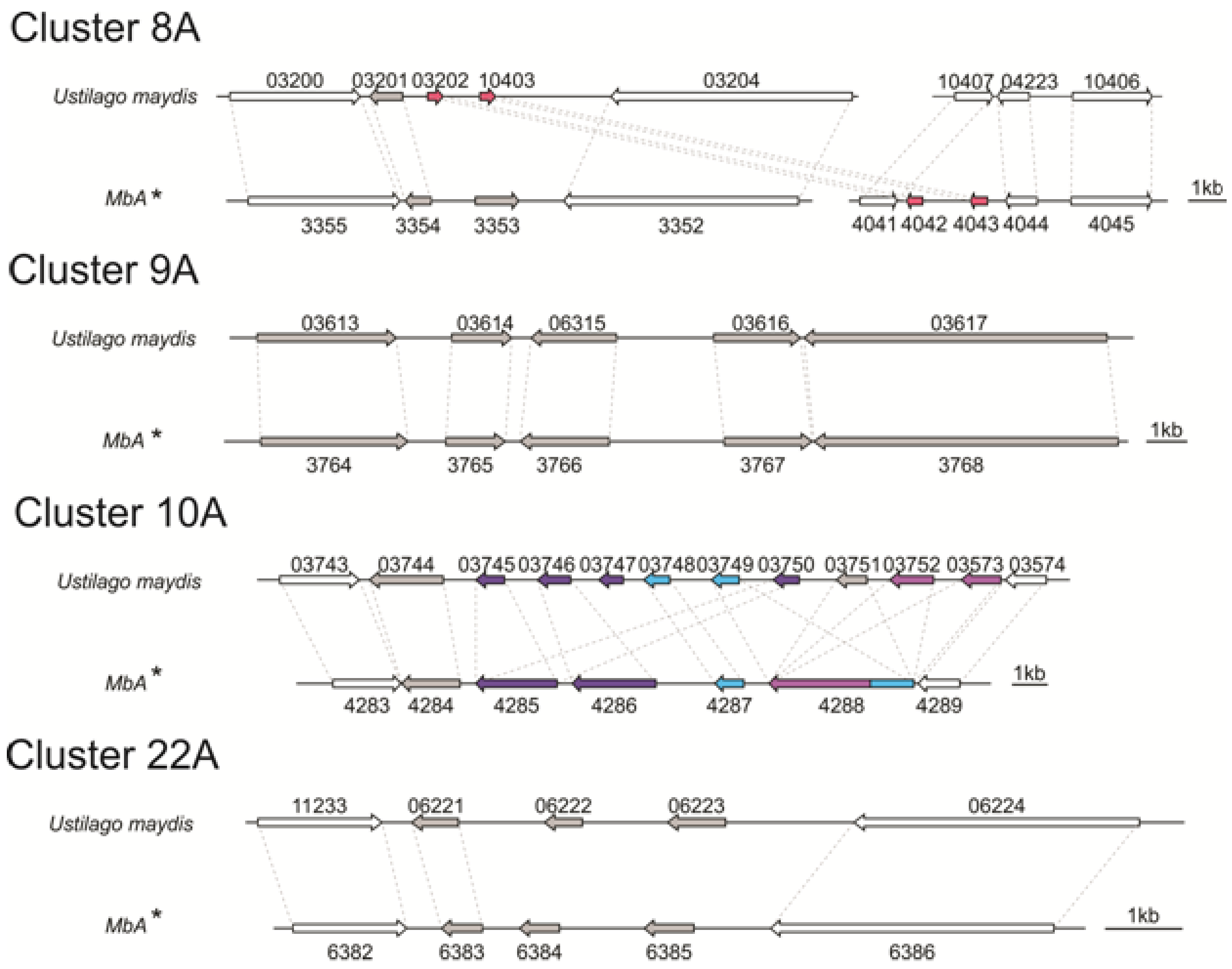


**Figure S6**: Comparison of known virulence clusters [4] between *U. maydis* and *MbA*. Numbers are gene numbers (*UMAG_NUMBER* for *U. maydis*; *gNUMBER* for *MbA).*

### Figure S7

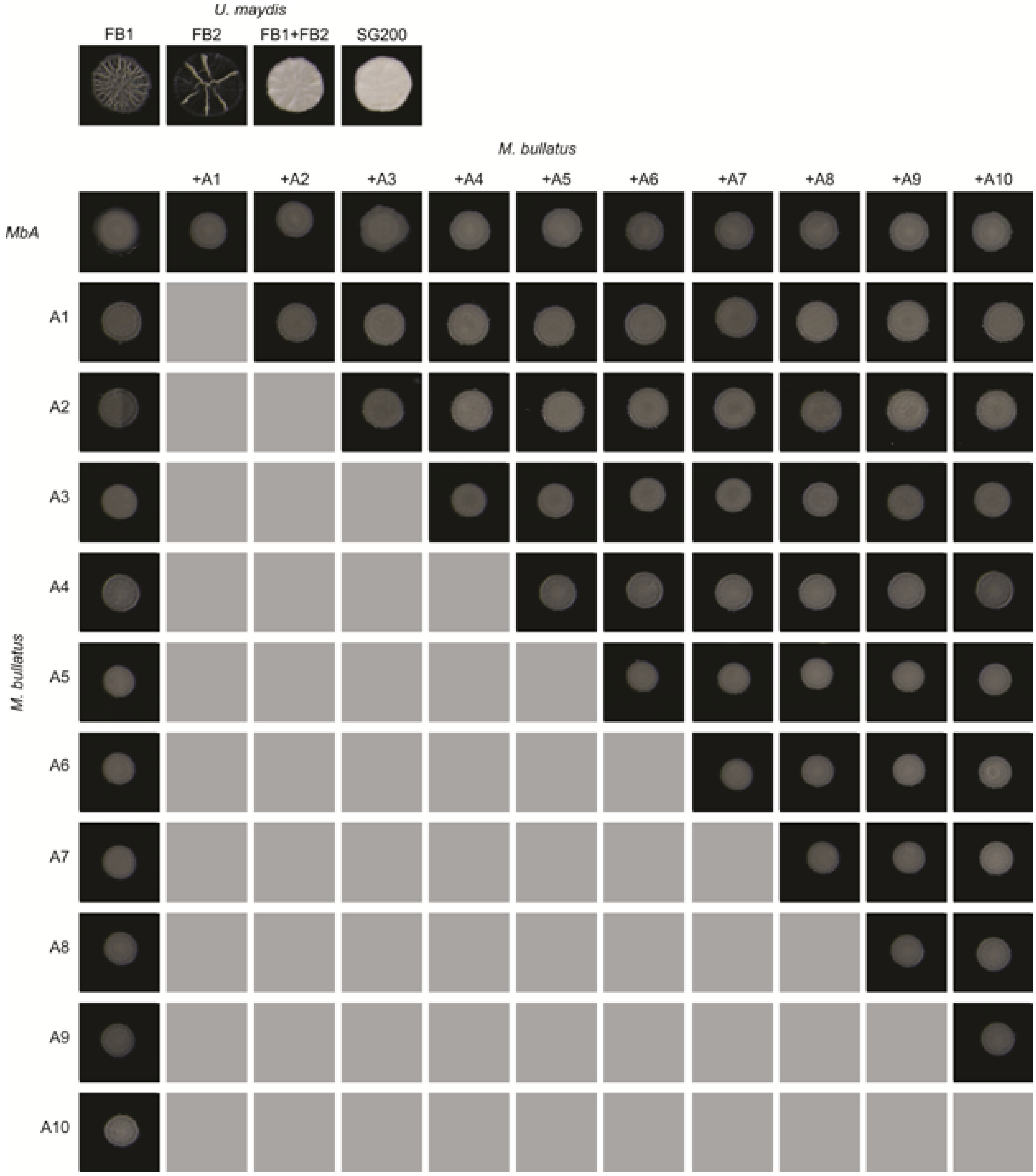


**Figure S7:** Mating assays of *MbA* and different *M. bullatus* isolates. Mixing of haploid *U. maydis* strains FB1 & FB2 and the solopathogenic strain SG200 served as a positive control for mating on charcoal plates, where filamentous growth is indicated by white, fluffy appearance of colonies. Haploid wild type strains without mating partner don't show a fluffy phenotype. For all combinations of *Moesziomyces* strains, no mating event resulting in filamentous growth on charcoal plates has been observed.

### Figure S8

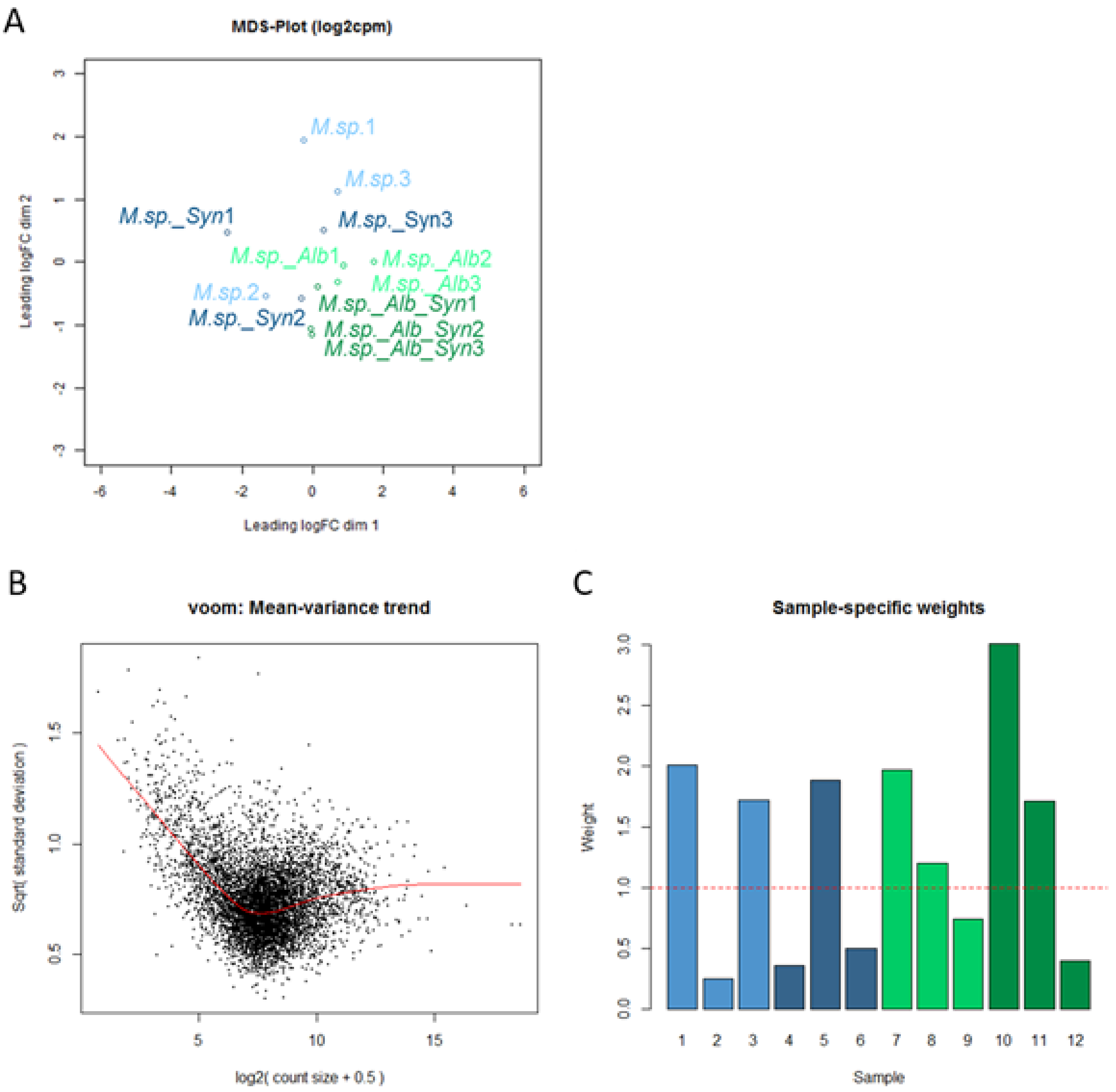


**Figure S8**: A) A) Multi-dimensional scaling plot (MDS) plot based on the interactions of *Moesziomyces* sp. (M.sp.) / *MbA* in response to the SynCom bacteria and *Albugo laibachii* in three biological replicates. The MDS plot shows *Albugo* and non-*Albugo* samples grouping together based on gene-level logCPM. (B) Voom mean-variance trend of the dataset where points represent genes, and (C) sample-specific weights obtained from the *limma-voom* function. Colours represent three replicates for each treatment. Light blue: *MbA* on plant; Dark blue: *MbA* on plant + SynCom; Light green: *MbA* on plant + Albugo; Dark green: *MbA* on plant + Albugo + SynCom.

**Figure S9:**

A

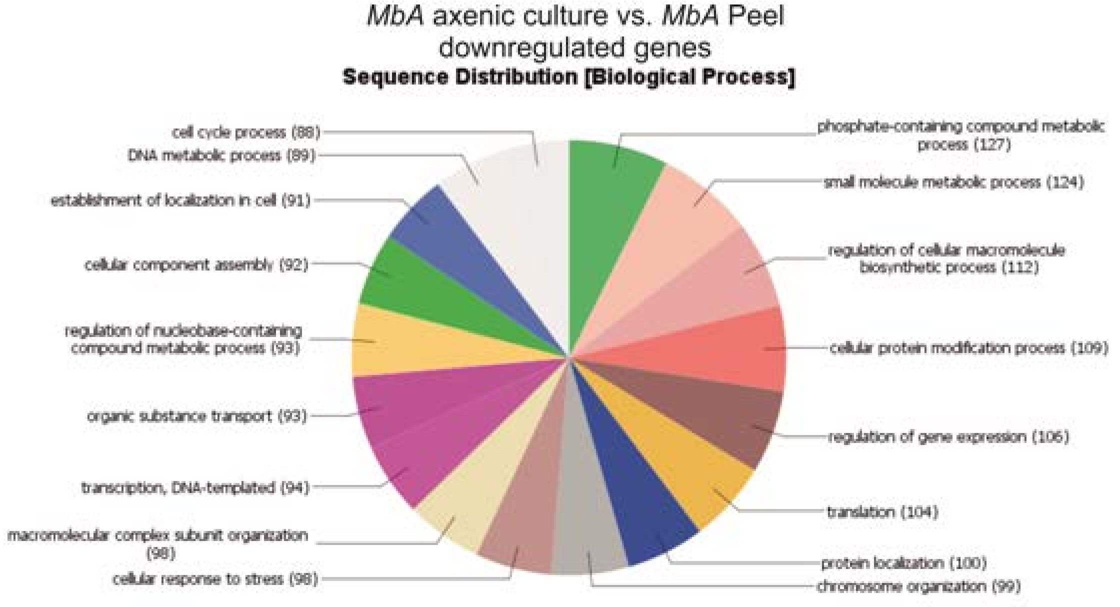


B

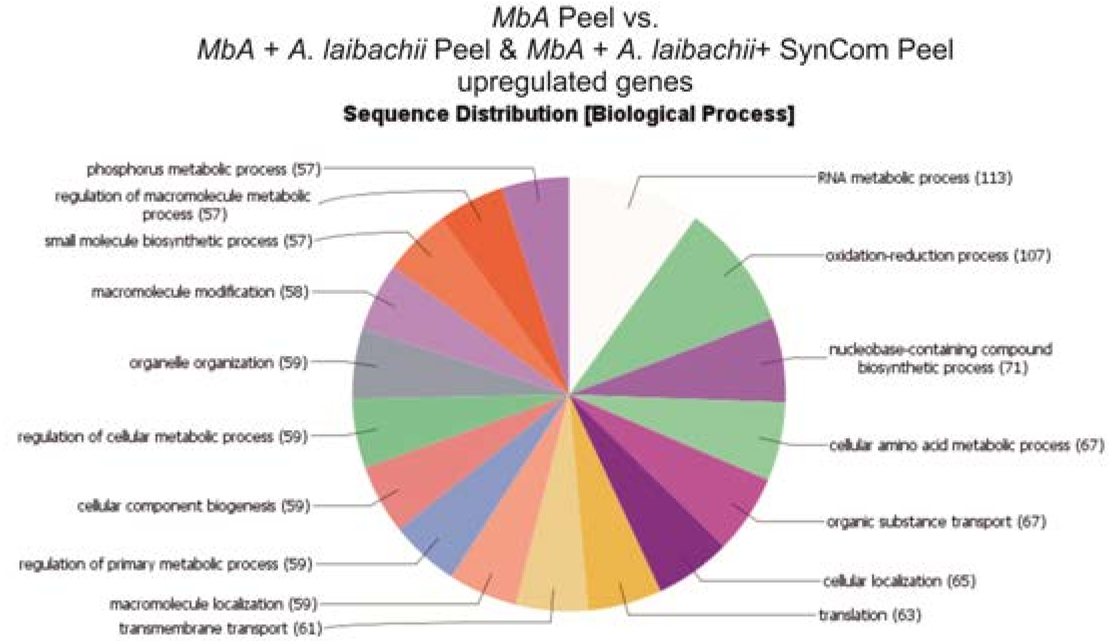


**Figure S9**: (A) Sequence distribution of Gene Ontology terms. 60% of all genes, downregulated in *MbA* on plant compared to axenic culture growth, can be assigned to GO-terms related to metabolism and cell cycle. (B) In contrast, presence of *A. laibachii* leads to transcriptional activation of metabolic processes. 52% of all GO-terms associated with genes upregulated in presence of *A. laibachii* are related to metabolic processes.

### Figure S10

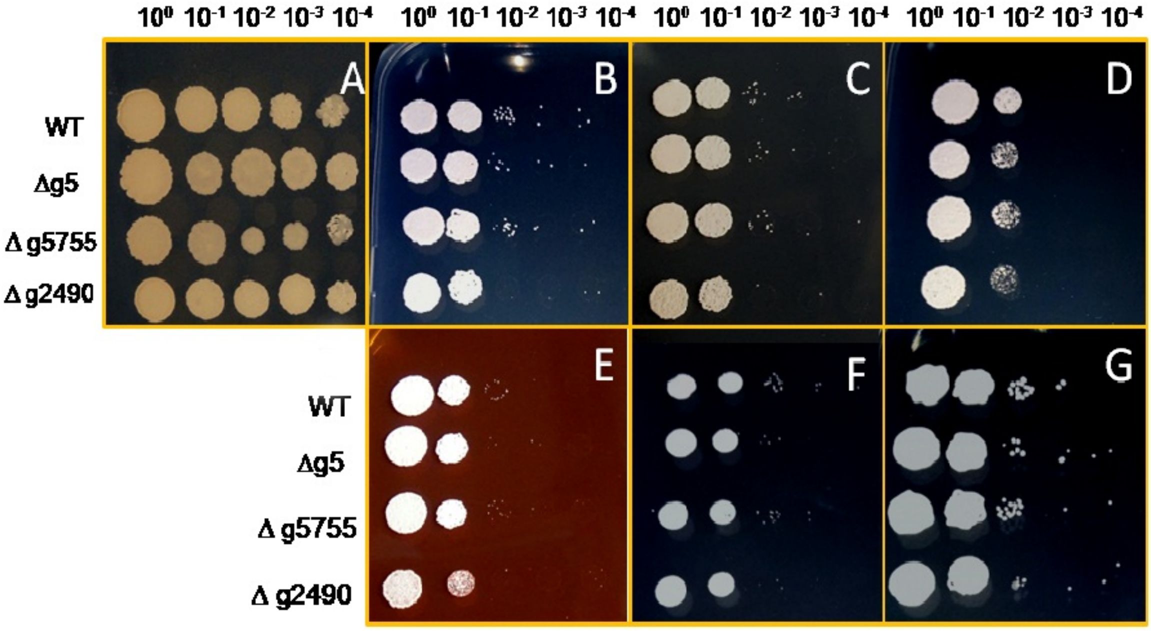


**Figure S10**: Stress assay of *MbA* wild type and knockout mutants of gene (g5, g5755 and g2490) respectively, on CM medium and 2% Glucose (A) with different conditions (B: 100 µg/ml Calcofluor; C: 150 µg/ml Calcoflour; D: 1 mMH2O2; E:45 µg/ml Congo-red; F: 1 M NaCl; G: 1 M Sorbitol). The strains were dropped on the CM plates containing different stress supplements in a dilution series from $10^0$ to $10^{-4}$

**Figure S11**

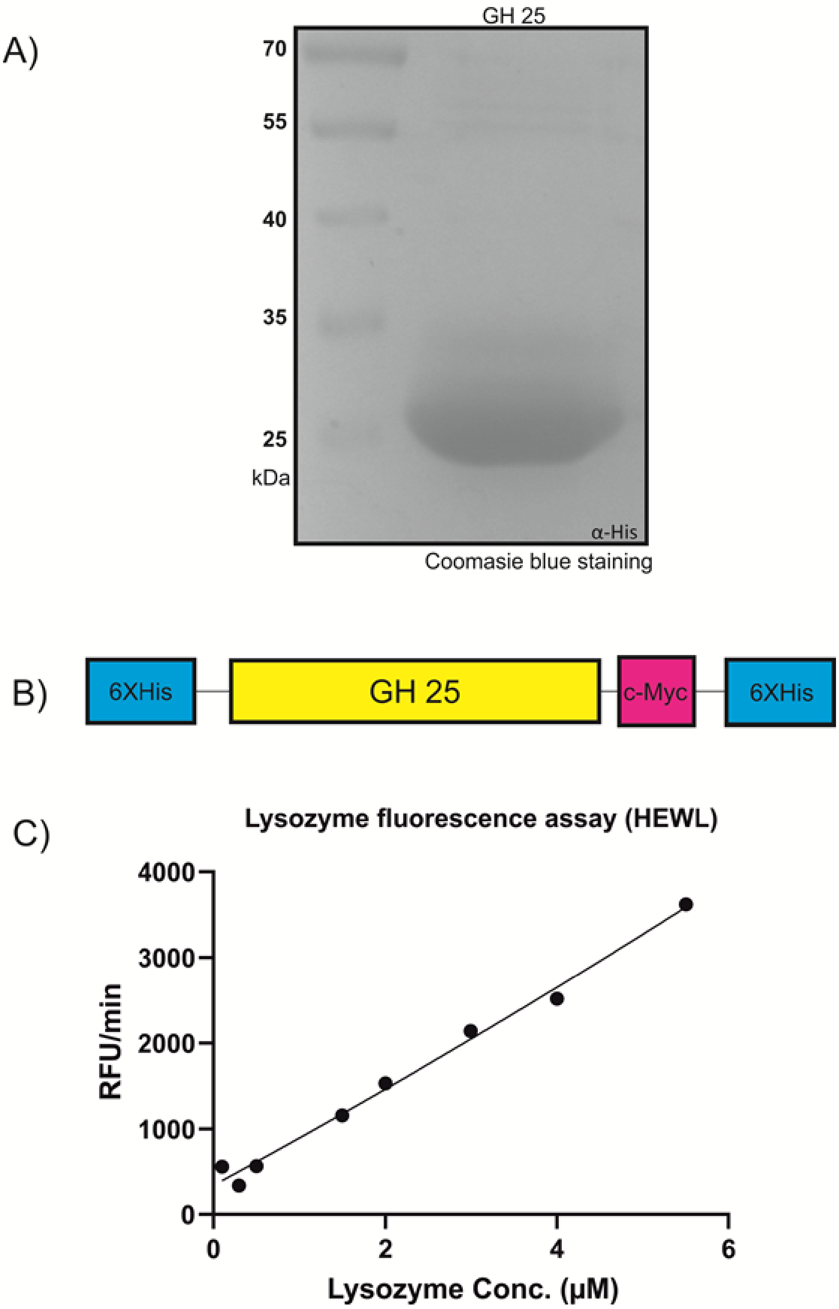


**Figure S11:** A) Recombinant MbA_GH25 was produced and purified using the *Pichia pastoris* protein expression system. The purified protein was loaded in a 12% SDS gel for visualization of an expected molecular weight of 27kDa for His-Tagged GH25. B) Schematic diagram of the recombinant construct, where the GH25 domain from *MBA* is tagged with an N-terminal polyhistidine Tag and a C-terminal peptide containing the c-myc epitope and a polyhistidine tag. C) Detection of lysozyme activity for Commercial Hen-egg white lysozyme (stock solution, 11µM) using the EnzChek[1] Lysozyme Assay Kit. The fluorescence was recorded every minute in a fluorescence microplate reader using excitation/emission of 485/530 nm in increasing concentrations from 0.1µM to 5.5 µM. Finally, Relative Fluorescence Unit (RFU)/ min was calculated for each concentration and plotted on the graph.

### Figure S12

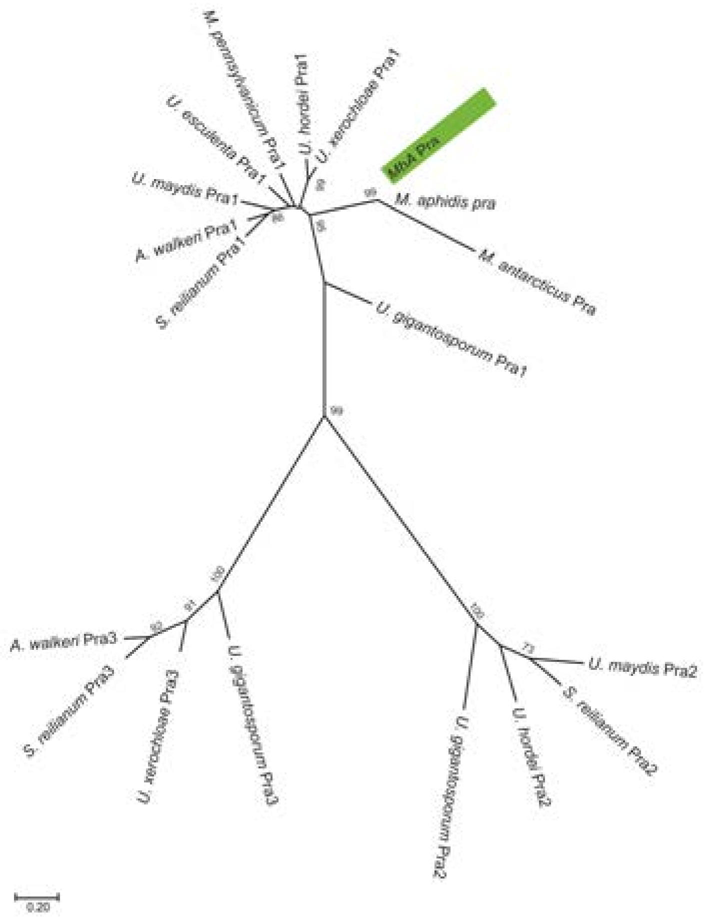


**Figure S12**: A molecular phylogenetic analysis by maximum likelihood method based on pheromone receptor protein sequences. MbA protein sequence clusters together with type 1 pheromone receptors of other Ustilaginomycetes.

### Figure S13:

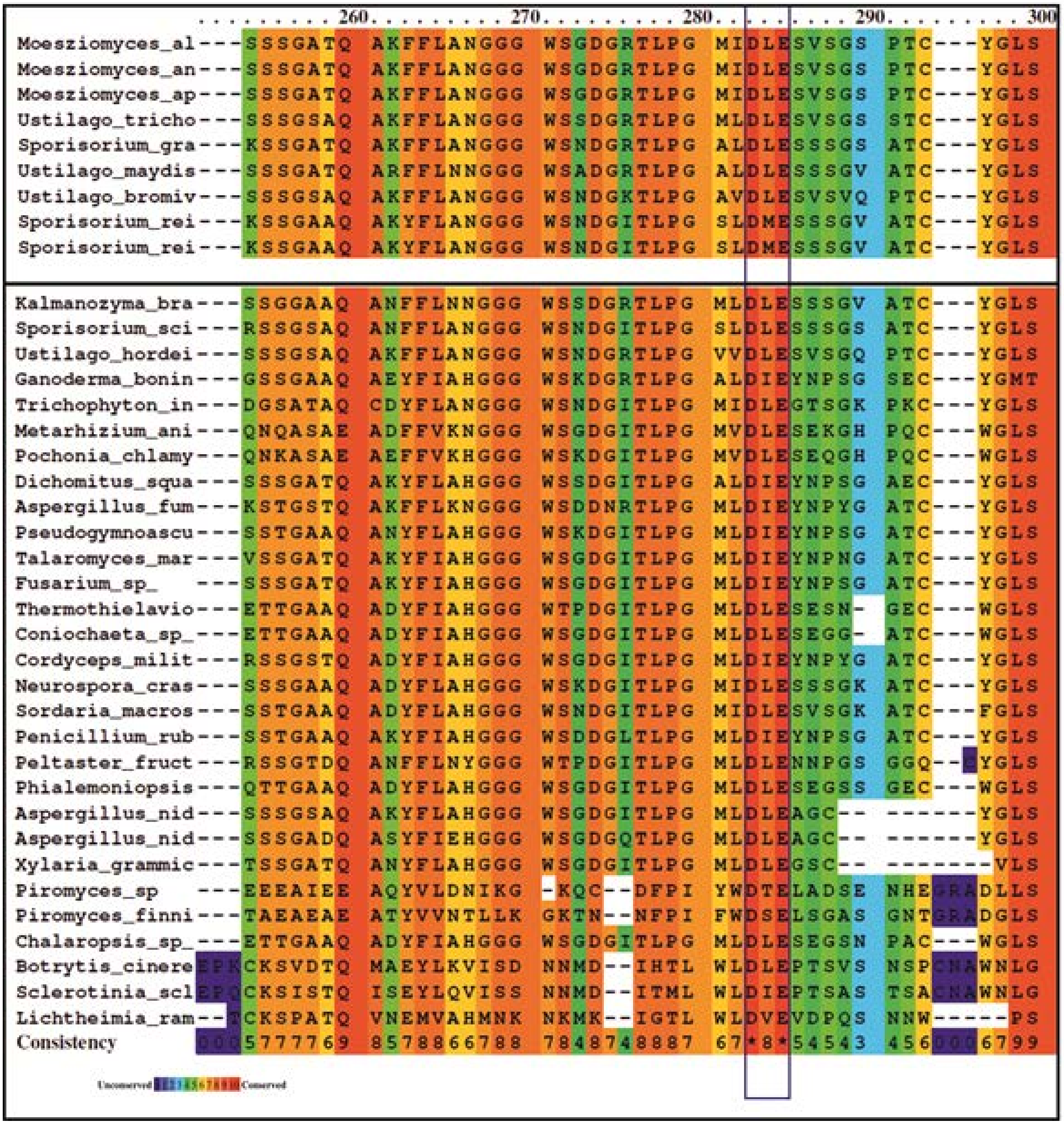


**Figure S13:** Amino acid alignment of GH25 sequences from different fungi (see attached list –'GH25 with accession number', for full length sequences). The protein sequences were obtained from the NCBI database. Alignment was achieved using the PRALINE multiple sequence alignment program with default parameters. The scoring scheme works from 0 for the least conserved alignment position, up to 10 (indicated by *) for the most conserved alignment position. A conserved active-site DxE motif has been predicted for glycoside hydrolase family 25. Sequences tested from different basiodiomycete, ascomycete and Chytrids, have the active site residue conserved (purple box).

### Figure S14:

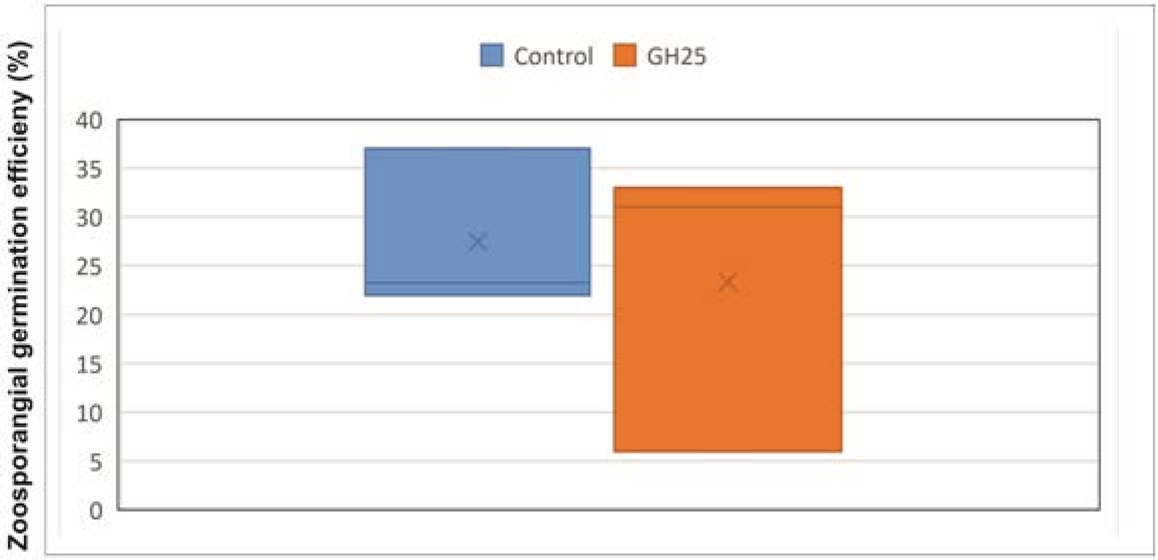


**Figure S14:** Boxplot-analysis of GH 25 treatment on *in vitro A. laibachii* zoosporangial germination in three biological replicates analyzing about 100 zoosporangial cells for each replicate. A p value of 0.3 was obtained for paired T-test using one-tailed distribution.
