## Supplementary material for "A fungal member of the *Arabidopsis thaliana* phyllosphere antagonizes *Albugo laibachii* via a secreted lysozyme": Table S1

| **Culture ID** | **Phyla** | **Family** | **Genus** | **Sampling location** | **Albugo infected Plant** | **Isolation data** | **Isolated on (media)** |
| --- | --- | --- | --- | --- | --- | --- | --- |
| 3 | *Proteobacteria* | *Pseudomonadaceae* | *Pseudomonas* | ERG | Y | 04.12.14 | Kings B |
| 4 | *Proteobacteria* | *Pseudomonadaceae* | *Pseudomonas* | ERG | Y | 04.12.14 | Kings B |
| 9 | *Bacterioidetes* | *Flavobacteriaceae* | *Flavobacterium* | ERG | N | 04.12.14 | Kings B |
| 10 | *Proteobacteria* | *Xanthomonadaceae* | *Xanthomonas* | ERG | N | 04.12.14 | Kings B |
| 18 | *Proteobacteria* | *Comamonadaceae* | *nd* | EY | N | 04.12.14 | Kings B |
| 20 | *Firmicutes* | *Listeriaceae* | *Brochothrix* | EY | N | 04.12.14 | Kings B |
| 24 | *Proteobacteria* | *Rhizobiaceae* | *nd* | JUG | N | 04.12.14 | Kings B |
| 25 | *Proteobacteria* | *Comamonadaceae* | *nd* | JUG | N | 04.12.14 | Kings B |
| 26 | *Proteobacteria* | *Pseudomonadaceae* | *Pseudomonas* | JUG | N | 04.12.14 | Kings B |
| 28 | *Bacterioidetes* | *Flavobacteriaceae* | *Flavobacterium* | JUG | Y | 04.12.14 | Kings B |
| 32 | *Proteobacteria* | *Moraxellaceae* | *Acinetobacter* | JUG | N | 04.12.14 | Kings B |
| 35 | *Actinobacteria* | *Microbacteriaceae* | *Microbacterium* | JUG | N | 04.12.14 | Kings B |
| 36 | *Actinobacteria* | *Microbacteriaceae* | *nd* | JUG | Y | 14.04.14 | Kings B |
| 38 | *Proteobacteria* | *Xanthomonadaceae* | *nd* | JUG | Y | 14.04.14 | Kings B |
| 40 | *Actinobacteria* | *Microbacteriaceae* | *nd* | JUG | Y | 14.04.14 | Kings B |
| 45 | *Firmicutes* | *Bacillaceae* | *Bacillus* | JUG | N | 14.04.14 | Kings B |
| 46 | *Actinobacteria* | *Streptomycetaceae* | *Streptomyces* | JUG | N | 14.04.14 | Kings B |
| 52 | *Actinobacteria* | *Micrococcaceae* | *Arthrobacter* | PFN | N | 14.04.14 | Kings B |
| 53 | *Actinobacteria* | *nd* | *nd* | PFN | N | 14.04.14 | Kings B |
| 54 | *Actinobacteria* | *Microbacteriaceae* | *Curtobacterium* | PFN | N | 14.04.14 | Kings B |
| 60 | *Proteobacteria* | *Pseudomonadaceae* | *Pseudomonas* | PFN | N | 14.04.14 | Kings B |
| 62 | *Actinobacteria* | *Microbacteriaceae* | *Microbacterium* | PFN | N | 04.12.14 | Kings B |
| 63 | *Firmicutes* | *Bacillaceae* | *Bacillus* | PFN | N | 04.12.14 | Kings B |
| 75 | *Proteobacteria* | *Pseudomonadaceae* | *Pseudomonas* | WH | N | 04.12.14 | Kings B |
| 76 | *Proteobacteria* | *Pseudomonadaceae* | *Pseudomonas* | WH | N | 04.12.14 | Kings B |
| 77 | *Proteobacteria* | *Xanthomonadaceae* | *nd* | WH | N | 04.12.14 | Kings B |
| 79 | *Proteobacteria* | *Burkholderiaceae* | *Pandoraea* | Al Nc14 culture | Y | - | LB |
| 80 | *Actinobacteria* | *Nocardiaceae* | *Rhodococcus* | Al Nc14 culture | Y | - | LB |
| 81 | *Proteobacteria* | *Xanthomonadaceae* | *Stenotrophomonas* | Al Nc14 culture | Y | - | LB |

**Table S1**: Members of the SynCom. The SynCom was compiled based on isolates that had been obtained from different natural sites (WH: Wendelsheim, PFN: Pfrondorf, JUG: Jugendhaus Hofgut Einsiedel EY: Eyach, ERG: Ergenzingen, Germany) and from *A. laibachii* isolate Nc14 spores. The isolates were chosen based on maximum diversity and repeated occurrence from different sampling sites. The column “Albugo infected plant” gives information if the plant that was sampled was infected by by A. laibachii or not. For all isolates partial 16S was sequenced using the common primer sets 799F and 1192R). Sequences were blasted to NCBI nr database and best hit listed if >98% identity.
