## Supplementary material for "A fungal member of the *Arabidopsis thaliana* phyllosphere antagonizes *Albugo laibachii* via a secreted lysozyme": Table S3

| Name | Composition |
| --- | --- |
| King’s B  medium | 20g Peptone (2%) (w/v), 1.5 (0.15%) (w/v) g K2HPO4 in 1 L double distilled (dd) water ~ pH adjusted to 7.2, 10ml (1%) (w/v) of Glycerol added, and autoclaved. Afterwards, 5 ml of (0.15%) 1M MgSO4 added. |
| YEPS light | 10 g (1% ) (w/v) Yeast-Extract, 0.4 g (0.4% ) (w/v) BactoTM –Peptone, 0.4 g (0.4% ) (w/v) Sucrose  mixed in 1L (dd) water and autoclaved |
| Regeneration agar | 15 g (1.5% ) (w/v) Bacto™ agar,182.2 g (18.22% ) (w/v) Sorbitol, 10 g (1% ) (w/v) Yeast extract, 0.4 g (0.4% ) (w/v) Bacto™-Peptone, and 0.4 g (0.4% ) (w/v) Sucrose mixed in 1L dd water and autoclaved. |
| Murashige skoog medium (1/2) | 1.1g MS salts (0.22%)(w/v), 4g (0.8%) (w/v) Bacto™ agar mixed in 500mL dd water and autoclaved. |
| CM medium | 2 g (0.25 %) (w/v) Casaminoacids (Difco)  0.8 g (0.1 %) (w/v) Yeast extract  8 ml (1 %) (v/v) Vitamine solution (Holliday ’74)  50 ml (6.25 %) (v/v) Salt solution (Holliday ’74)  0.4 g (0.05 %) (w/v) DNA degradation free (Sigma)  1.2 g (0.15 %) (w/v) NH4NO3  10 ml (1 %) (v/v) 1 M Tris-HCl pH 8.0  16 g (2 %) (w/v) Agar  The components are mixed in 784 ml ddH2O, and autoclaved. After autoclaving, 16 ml 50 % Glucose (sterile filtrated) is added to get a final concentration of 2% glucose.  Stress conditions  CM + 2 % Glucose (Control)  CM + 2 % Glucose + 100 μg/ml Calcofluor (cell wall stress)  CM + 2 % Glucose + 150 μg/ml Calcoflour (cell wall stress)  CM + 2 % Glucose + 1 mM H2O2 (oxidative stress)  CM + 2 % Glucose + 1.5 mM H2O2 (oxidative stress)  CM + 2 % Glucose + 45 μg/ml Congored (cell wall stress)  CM + 2 % Glucose + 1 M NaCl (osomotic stress)  CM + 2 % Glucose + 1 M Sorbitol (osmotic stress) |
| PD Agar | 2.4% (w/v) Potato dextrose broth, 2% (w/v) Bacto-Agar |
| PD charcoal medium | 2.4 % (w/v) Potato-Dextrose Broth, 2 % (w/v) Bacto Agar 1% (w/v) Charcoal (Sigma), 1% 1(M) Tris-HCl pH 8.0 |
| SCS | 20 mM Na-citrate(pH 5), 1 M Sorbitol ~ Sterile-filtered in ddH2O |
| SCS/Glucanex | 3ml SCS with 60 mg (2 %) Glucanex |
| STC | 10 mM Tris-HCl, pH 7.5 ,100 mM CaCl2, 1 M Sorbitol  Sterile-filtered in ddH2O |
